# The role of microglial LRRK2 in manganese-induced inflammatory neurotoxicity via NLRP3 inflammasome and RAB10-mediated autophagy dysfunction

**DOI:** 10.1101/2023.04.03.535418

**Authors:** Edward Pajarillo, Sang Hoon Kim, Alexis Digman, Matthew Dutton, Deok-Soo Son, Michael Aschner, Eunsook Lee

**Affiliations:** Department of Pharmaceutical Science, College of Pharmacy and Pharmaceutical Sciences, Florida A&M University, Tallahassee, FL, USA 32307; Department of Biochemistry and Cancer Biology, Meharry Medical College, Nashville, TN, USA, 37208; Department of Molecular Pharmacology, Albert Einstein College of Medicine, Bronx, New York, New York, USA, 10461

**Keywords:** manganese, microglia, inflammation, LRRK2, G2019S, NLRP3 inflammasome, RAB10, autophagy, lysosome

## Abstract

Chronic exposure to manganese (Mn) can lead to manganism, a neurological disorder sharing common symptoms with Parkinson’s disease (PD). Studies have shown that Mn can increase the expression and activity of leucine-rich repeat kinase 2 (LRRK2), leading to inflammation and toxicity in microglia. LRRK2 G2019S mutation also elevates LRRK2 kinase activity. Thus, we tested if Mn-increased microglial LRRK2 kinase is responsible for Mn-induced toxicity, and exacerbated by G2019S mutation, using WT and LRRK2 G2019S knock-in mice, and BV2 microglia. Mn (30 mg/kg, nostril instillation, daily for 3 weeks) caused motor deficits, cognitive impairments, and dopaminergic dysfunction in WT mice, which were exacerbated in G2019S mice. Mn induced proapoptotic Bax, NLRP3 inflammasome, IL-1β and TNF-α in the striatum and midbrain of WT mice, and these effects were exacerbated in G2019S mice. BV2 microglia were transfected with human LRRK2 WT or G2019S, followed by Mn (250 μM) exposure to better characterize its mechanistic action. Mn increased TNF-α, IL-1β, and NLRP3 inflammasome activation in BV2 cells expressing WT LRRK2, which was exacerbated in G2019S-expressing cells, while pharmacological inhibition of LRRK2 mitigated these effects in both genotypes. Moreover, the media from Mn-treated BV2 microglia expressing G2019S caused greater toxicity to cath.a-differentiated (CAD) neuronal cells compared to media from microglia expressing WT. Mn-LRRK2 activated RAB10, which was exacerbated in G2019S. RAB10 played a critical role in LRRK2-mediated Mn toxicity by dysregulating the autophagy-lysosome pathway, and NLRP3 inflammasome in microglia. Our novel findings suggest that microglial LRRK2 via RAB10 plays a critical role in Mn-induced neuroinflammation.

## Introduction

Chronic exposure to elevated levels of manganese (Mn) from both occupational and environmental sources can lead to its accumulation in various brain regions, particularly the basal ganglia (1,2). This leads to a neurological disorder referred to as manganism, manifesting as Parkinson’s disease (PD)-like pathological symptoms, such as tremor, dystonia, abnormal gait, and bradykinesia in Mn-exposed welders and adults (3–5). In children, high levels of Mn have been linked to motor and behavioral deficits. Experimental rodent models exposed to high levels of Mn have also shown movement deficits and impaired motor coordination with concomitant nigrostriatal dopaminergic dysfunction in the mouse brain (1,6). In addition to motor deficits, Mn toxicity has been shown to cause cognitive dysfunction, including speech impediment and memory impairment (7,8). Studies have revealed that high Mn exposure levels in children and adults are linked to cognitive and memory deficits (7,9), and Mn induced hippocampal dysfunction and memory impairment in experimental animal models (10,11). This growing evidence underscores Mn’s crucial role in causing motor deficits and cognitive impairment, and the urgency for understanding the underlying mechanisms of Mn-induced neurotoxicity as to enable development of efficacious therapeutics.

The precise cellular and molecular mechanisms of Mn-induced neurotoxicity are not fully understood, but studies have shown that Mn can induce oxidative stress, inflammation, apoptosis, excitotoxicity, and autophagy dysregulation in various neural cell types (for review, see (12)). Among these mechanisms, inflammation induced by glial cells, particularly microglia, is considered critical for Mn-induced neuroinflammation (12–14). Mn increases the production of proinflammatory cytokines, such as tumor necrosis factor (TNF)-α, interleukin (IL)-1β, IL-6, interferon γ, and chemokines in microglia (13–17), which are released and damaging adjacent cells including neurons by paracrine and exosome modes, causing inflammatory neuronal dysfunction.

The leucine-rich repeat kinase 2 (LRRK2) is a protein kinase, also known as dardarin encoded by PARK8 gene, containing multiple functional domains, including GTP-binding and kinase domains (for review, see (18)). Among many LRRK2 mutations identified, LRRK2 mutation on G2019S is most common and associated with clinical cases of familial and idiopathic late-onset PD (18,19). G2019S mutation is also related to dementia with Lewy bodies which have shown to contain aggregates of dysfunctional proteins, including LRRK2 (20). LRRK2 regulates vesicle trafficking, autophagy-lysosomal pathway, and neurite development (for review, see (18)). Intriguingly, mutations in LRRK2 at G2019S, R1441C/G/H, and I2020T are known to induce gain-of-function of LRRK2 kinase activity. Several studies have implicated gain-of-function of LRRK2 activity in the G2019S mutant. G2019S altered LRRK2’s protein structure, potentially affecting binding and accessibility of the kinase domain (21,22). LRRK2 mutation at R1441H, which is located close to the GTPase domain could increase GTPase activity, which in turn, enhance its kinase activity (23). Regardless of mechanisms, ultimate LRRK2’s hyper-kinase activity will likely result in abnormally increased phosphorylation of LRRK2 substrates, including several RAB proteins (24). RAB proteins are small GTPases involved in regulating membrane trafficking, centrosome structure, and lysosomal function (for review, see (25)), and dysregulation of RAB proteins has been associated with PD pathogenesis (for review, see (25)). Among the RAB proteins, RAB10 is known to play a critical role in the endoplasmic reticulum, endosomes, dendritic growth, and lysosomal function (26–28), and its increased phosphorylation at threonine 73 (pT73) has been implicated in LRRK2 G2019S mutation-related PD and LRRK2 hyper kinase activity (26). Elevated RAB10 phosphorylation was also observed in PD patients with LRRK2 mutation sites other than G2019S (29), indicating that RAB10 phosphorylation could be a relevant biomarker of LRRK2 activity and dysfunction in PD patients (25).

Gene-environment interactions have been proposed as potential contributors to the varied clinical manifestations of idiopathic PD, because not all carriers of PD-related LRRK2 mutations exhibit PD pathology (20). In addition, studies have shown that LRRK2 played a role in Mn-induced cytotoxicity and inflammation in microglial cells and mice (13,30). Mn increased LRRK2 expression, its kinase activity, and proinflammatory cytokines TNF-α and IL-1β with concomitant impairment of autophagy (30). Mn-induced apoptosis and cytokine production were mediated by increased LRRK2 kinase activity in microglia as pharmacological inhibition of LRRK2 with GSK2578215A and MLi-2 abolished Mn-induced oxidative stress, apoptosis, and proinflammatory TNF-α production in HMC3 human microglial cells (13). Despite these findings, the mechanism of Mn-induced toxicity via LRRK2 in microglia has yet to be established. Studies have shown that Mn-induced LRRK2 hyper kinase activity could modulate LRRK2’s downstream targets, such as RAB, MAPK p38, and ERK (13). Mn also dysregulated autophagy proteins such as ATG5 and Beclin 1, while inhibition of LRRK2 with LRRK2 siRNA and inhibitor LRRK2-IN-1 abolished these Mn effects in BV2 microglia (30). Studies have also shown that Mn disrupted Golgi trafficking and distribution in HeLa cells, partially via impairment RAB-dependent signaling (31). Thus, its critical to investigate whether LRRK2-RAB pathway is involved in Mn-induced impairment in autophagy and lysosomal function, leading to microglial inflammation and adjacent neuronal injury.

In this study, we examined the Mn’s toxic effects in the mouse midbrain and striatum where nigrostriatal dopaminergic neurons are innervated and are known targets of Mn’s neurotoxicity in both the LRRK2 wild type (WT) and G2019S knock-in (KI) genotypes. We tested if LRRK2 kinase was involved in Mn-induced neurotoxicity in the nigrostriatal region by inflammatory neurotoxicity via increase of TNF-α and NLRP3/IL-1β inflammasome, and if these Mn effects are exacerbated in G2019S mice due to its hyper LRRK2 kinase. We also carried out an in vitro study in BV2 cells expressing LRRK2 WT and G2019S to investigate and gain more insight on the mechanistic role of microglial LRRK2 in mediating Mn-induced neurotoxicity and neuroinflammation.

Our findings reveal that Mn induced inflammatory neurotoxicity in the nigrostriatal region of the mouse brain of WT mice, which were exacerbated in G2019S mice characterized by increased dopaminergic dysfunction, motor deficits, and cognitive impairment, along with molecular alterations in LRRK2, tyrosine hydroxylase (TH), TNF-α, and IL-1β. These effects appeared to be mediated at least in part through the microglial LRRK2-RAB10-NLRP3 pathway as evidenced by results derived from in vitro BV2 microglial cultures.

## Results

### Mn-induced motor deficits in WT mice were exacerbated in LRRK2 G2019S mice

We have previously shown that Mn increased LRRK2 expression and kinase activity in microglia and the mouse brain, contributing to inflammatory toxicity (13,30). LRRK2 G2019S mutation also increased LRRK2 kinase activity (32,33), suggesting that elevated kinase activity of LRRK2 is a common factor in both Mn and G2019S pathology. Although G2019S is associated with an increased risk of familial as well as idiopathic PD (20,34,35), its penetrance of PD is incomplete (36), and the risk of developing PD increases with age (37). Thus, we investigated if Mn contributes to pathological symptoms in G2019S mice using 8-week-old WT and G2019S KI mice. The results showed that three weeks of Mn exposure (30 mg/kg) via nostril instillation significantly decreased locomotor activities, such as total distance traveled, walking speed, and vertical activities in the WT mice. There was no significant difference in locomotor activity between WT and G2019S mice without Mn in this age of mice (8-week-old), suggesting that the low penetrance of PD in G2019S mice. However, Mn-induced motor deficits were higher in G2019S mice compared to those in WT mice (Fig. 1A-D). Mn also impaired motor coordination, reflecting the ability of mice to stay on the rotarod, and this Mn effect was exacerbated in G2019S mice (Fig. 1E). In the absence of Mn, the rotarod performance of G2019S mice was like that of WT mice.

**Figure 1.**
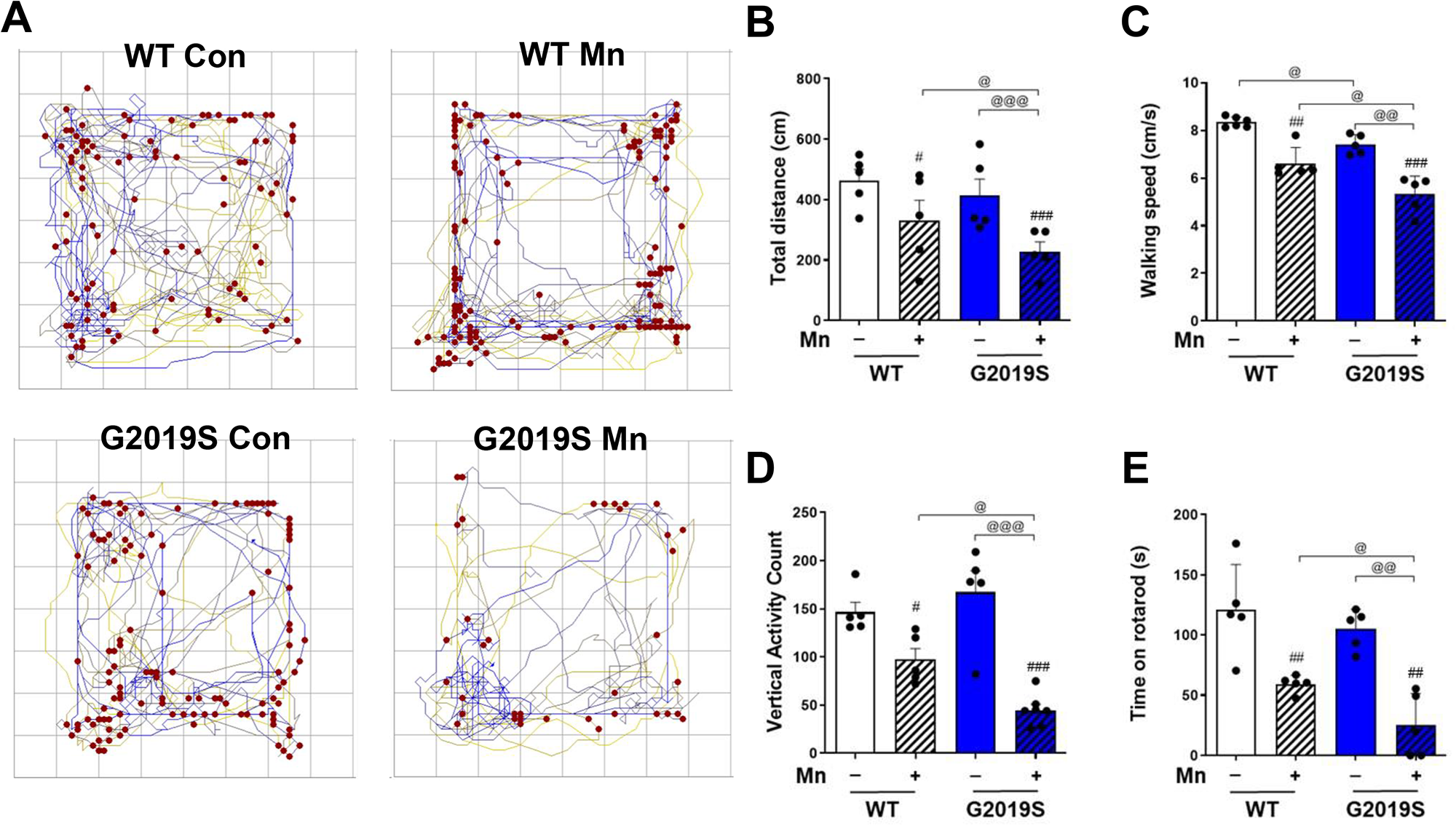
Mn-induced movement impairment and locomotor deficits is further worsened in mice with enhanced LRRK2 kinase activity (LRRK2 G2019S KI). (A-D) After Mn exposure (MnCl_2_, 30 mg/kg, intranasal instillation, daily for 3 weeks), locomotor activity was measured by open-field traces (A), total distance traveled (B), walking speed (C), and vertical activity count (D). The mouse’s movement is depicted by traces and red dots indicate the location of vertical activity in the open-field arena. (E) Motor coordination is measured by fall latency as time spent on the rotating rod. ^#^p < 0.05, ^##^p < 0.01, ^###^p < 0.001, compared with the controls; ^@^p < 0.05, ^@@^p < 0.01, ^@@@^p < 0.01, compared with each other (two-way ANOVA followed by Tukey’s post hoc test; n = 5). Data are expressed as mean ± SD.

### Mn-induced cognitive impairment is exacerbated by LRRK2 G2019S mutation in mice

Cognitive impairment has been well established in Mn toxicity (38,39), and LRRK2 G2019S mutation has also been associated with dementia (40). To determine whether Mn-induced cognitive dysfunction is altered by LRRK2 G2019S mutation, we performed a novel object (NO) recognition test, which reflects the mouse’s time spent exploring and interacting with NO in WT and G2019S mice (41). Our results showed that, while LRRK2 G2019S mutation did not affect the mice’s ability to recognize NO, Mn exposure decreased NO recognition in WT mice, an effect which was exacerbated in G2019S mice (Fig. 2A-C). The latter exhibited reduced time spent exploring and interacting with NO (Fig. 2B) and lower scores for their ability to differentiate NO from the familiar object (FO) compared to the WT group (Fig. 2C).

**Figure 2.**
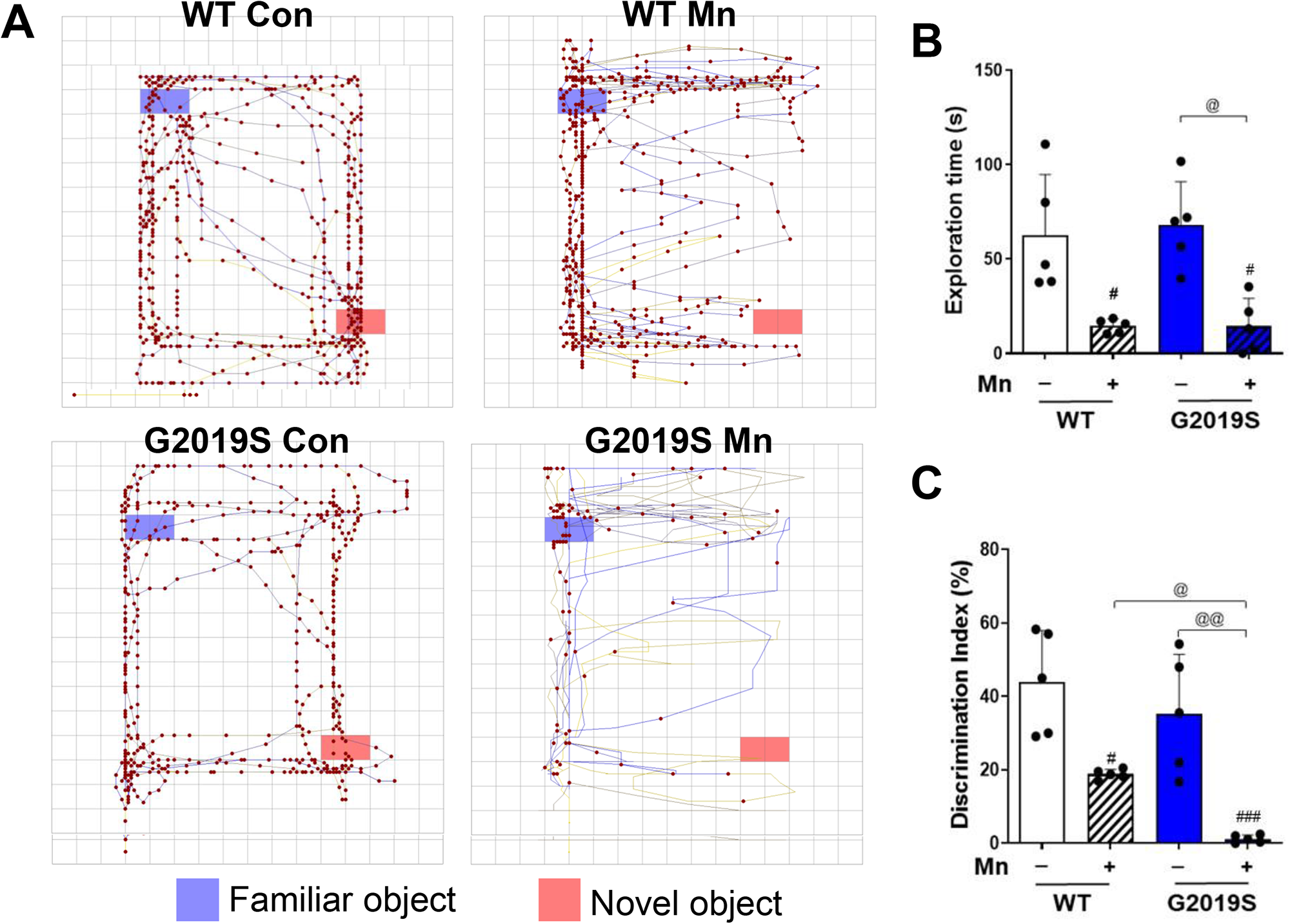
Mn-induced cognitive impairment is exacerbated in LRRK2 G2019S mice. (A-C) After Mn exposure (MnCl_2_, 30 mg/kg, intranasal instillation, daily for 3 weeks), NO recognition was evaluated as described in the methods section. (A) NO recognition in mice was assessed by comparing the time spent on the NO (bottom-right) compared to the FO (top-left). Traces in the arena depict the mouse’s movement and interaction with FO and NO over a 10-min period. Red dots on the FO and NO shows a point of exploration with an object. (B-C) The time spent in each object in the open-field arena was used to calculate the NO exploration time (B) and discrimination index (C). ^#^p < 0.05, ^###^p < 0.001, compared with the controls; ^@^p < 0.05, ^@@^p < 0.01, compared with each other (two-way ANOVA followed by Tukey’s post hoc test; n = 5). Data are expressed as mean ± SD.

### Mn-induced dopaminergic neuronal dysregulation is exacerbated in LRRK2 G2019S mice

Mechanisms underlying Mn-induced motor deficits are not fully established, but Mn impairs dopaminergic neuronal functions (1,42). Decreased expression of TH, a rate-limiting enzyme for dopamine synthesis, in the nigrostriatal region has been verified (2,42,43). To determine if Mn-induced dopaminergic dysfunction is worsened in LRRK2 G2019S mice, we measured extracellular dopamine levels, a readout of dopamine release, in the mouse striatum by microdialysis, and TH protein levels in the striatum and substantia nigra (midbrain) where dopaminergic neurons are innervated. Acute Mn (50 mg/kg, i.p., 1 h) treatment significantly reduced striatal dopamine levels stimulated by 100 mM KCl in WT mice, and this effect was further reduced in G2019S mice (Fig. 3A, B). Without Mn treatment, G2019S mice did not reduce striatal dopamine levels significantly.

**Figure 3.**
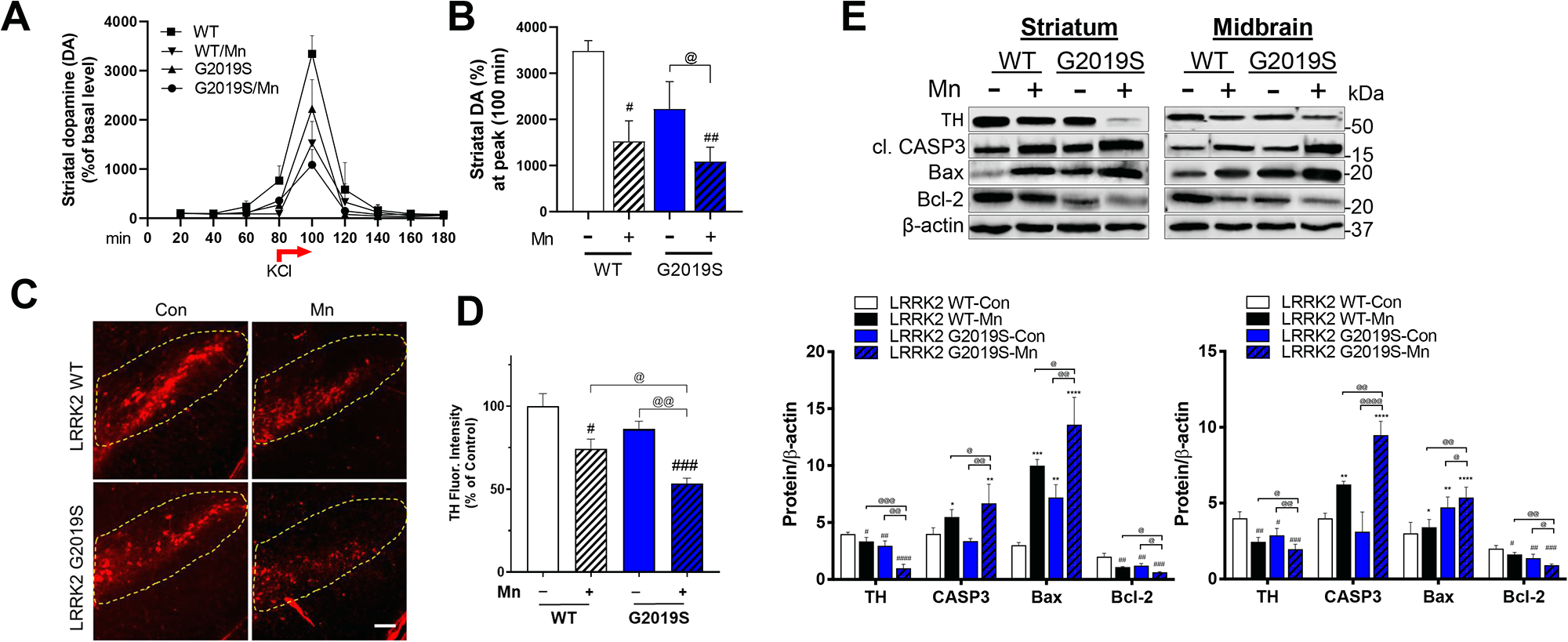
Mn-induced dysfunction of nigrostriatal dopaminergic pathway is greater in LRRK2 G2019S than in WT mice. (A-B) After acute Mn exposure, extracellular dopamine levels in the mouse striatum were measured by microdialysis and HPLC-ECD, as described in the Methods section. (A) Extracellular dopamine release was stimulated after 100 mM KCl, for dopamine release for 20 min between 80-100 min intervals. (B) Striatal dopamine levels were compared among different groups. (C) After Mn exposure (MnCl_2_, 30 mg/kg, intranasal instillation, daily for 3 weeks), coronal sections of the substantia nigra were immunostained for TH proteins by IHC as described in the Methods section. (D) TH fluorescence intensities were compared among different treatment groups. (E) After Mn exposure (MnCl_2_, 30 mg/kg, intranasal instillation, daily for 3 weeks), protein levels of TH, cleaved caspase-3, Bcl-2, and Bax were measured in the striatum and midbrain. β-actin was used as a loading control for protein. *p < 0.05, **p < 0.01, ***p < 0.001, ****p < 0.0001, ^#^p < 0.05, ^##^p < 0.01, ^###^p < 0.001, ^####^p < 0.0001, compared with the controls; ^@^p < 0.05, ^@@^p < 0.01, ^@@@^p < 0.001, ^@@@@^p < 0.0001, compared with each other (two-way ANOVA followed by Tukey’s post hoc test; n =3-5). Data are expressed as mean ± SD.

G2019S reduced TH protein levels in the striatum and midbrain compared to WT even without Mn treatment (Fig. 3E), but this reduction was not correlated with decreased motor function (Fig. 1). Mn effects on TH protein expression by immunofluorescence signals in the brain sections from substantia nigra showed similar reduction in WT mice, which was exacerbated in G2019S mice (Fig. 3C, D). We also tested apoptotic proteins, such as B-cell lymphoma 2 (Bcl-2), Bcl-2 associated protein X (Bax), and cleaved caspase-3 (CASP3). The results showed that Mn decreased protein levels of antiapoptotic Bcl-2 and increased proapoptotic Bax and cleaved CASP3 in the striatum and midbrain of WT mice (Fig. 3E), effects which were further exacerbated in G2019S mice. These results indicate that LRRK2 plays a role in Mn-induced neurotoxicity in the nigrostriatal dopaminergic pathway, at least in part by increasing proapoptotic proteins and decreasing TH.

### Mn increased LRRK2 expression and its kinase function along with increasing microglial inflammation and proinflammatory TNF-α production in the mouse brain

Given that Mn increased LRRK2 expression in the mouse brain and microglia (30), as well as enhanced LRRK2 kinase activity by autophosphorylation of LRRK2 at S1292 (13), we tested if these effects were modulated in G2019S which have inherently higher LRRK2 kinase activity compared to WT (33,44), focusing on microglial changes. Results showed that Mn increased LRRK2 protein levels in the striatum and midbrain of WT and further increased in G2019S mice (Fig. 4A). These effects on LRRK2 protein levels in the substantia nigra were accompanied by an increase in the microglial activation as reflected by higher fluorescence of the microglial marker, Iba1 (Fig. 4B). Furthermore, Mn increased colocalization between LRRK2 and Iba1 in the mouse substantia nigra, indicating that this Mn-induced increase in LRRK2 protein expression inherent to microglial function.

**Figure 4.**
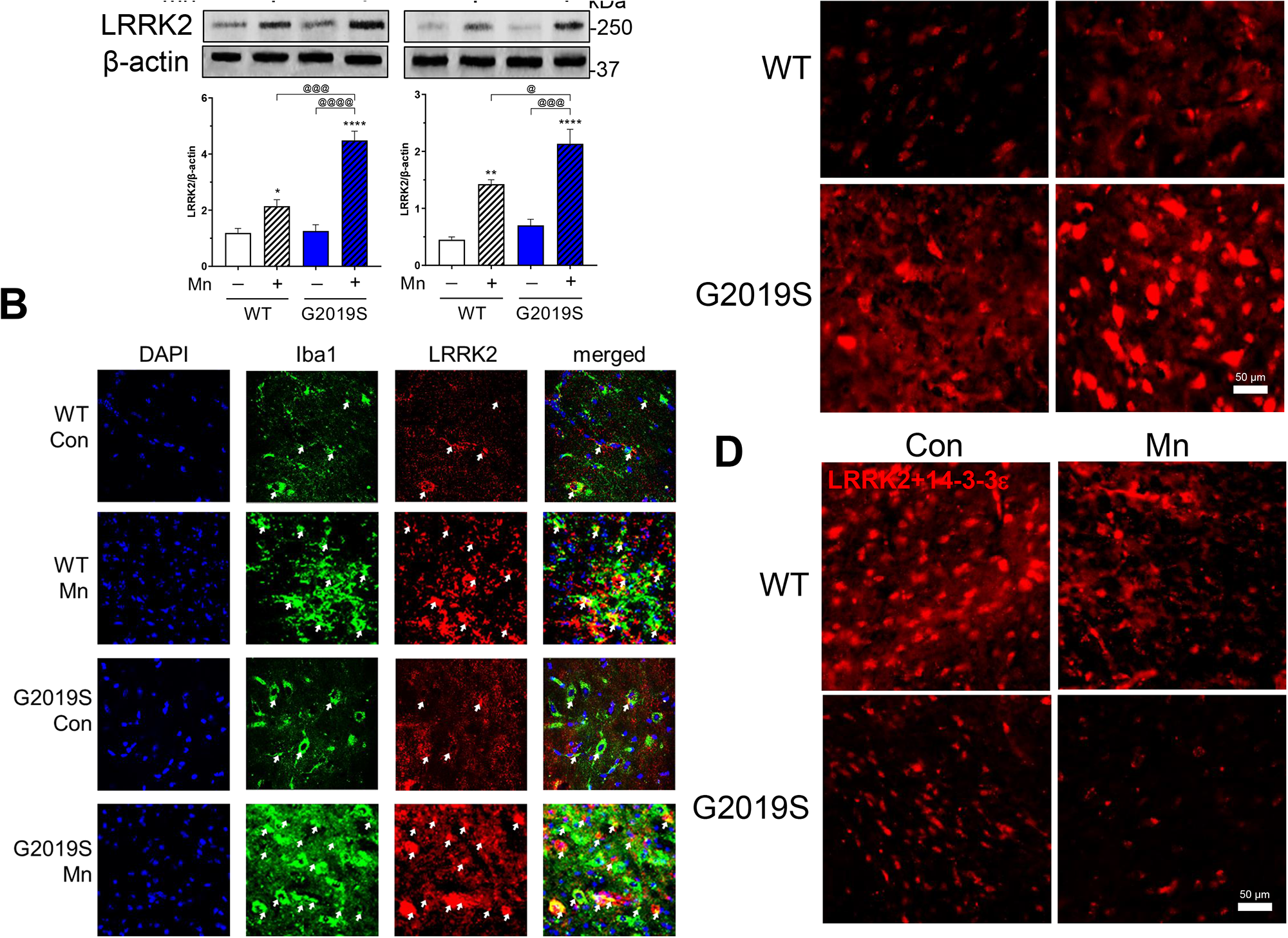
Mn increased microglial LRRK2 expression and LRRK2 activity in the mouse nigrostriatal region. (A-B) After Mn exposure (MnCl_2_, 30 mg/kg, intranasal instillation, daily for 3 weeks), LRRK2 protein levels were analyzed in the striatum and midbrain regions of mouse brains by western blotting (A) and IHC of the substantia nigra region (B). β-actin was used as a loading control. (A) Protein levels of LRRK2 in striatum and midbrain of Mn-treated WT and G2019S mice. (B) Expression and colocalization of LRRK2 and Iba1, a microglial marker, were visualized with red and green fluorescence signals, respectively. (C-D) PLA for protein-protein interactions of LRRK2 with RAB10 (C) and LRRK2 with 14-3-3ξ (D) in the substantia nigra was visualized with red fluorescence signals. β-actin was used as a loading control for protein. *p < 0.05, **p < 0.01, ****p < 0.0001, compared with the controls; ^@^p < 0.05, ^@@@^p < 0.001, ^@@@@^p < 0.0001, compared with each other (two-way ANOVA followed by Tukey’s post hoc test; n = 3). Data are expressed as mean ± SD.

Next, we tested if Mn-induced increase of LRRK2 results in activation of LRRK2 kinase substrates, particularly RAB10. We assessed the protein-protein interaction between LRRK2 and RAB10 or 14-3-3, a regulatory protein of LRRK2 function and may play a role in PD (45,46). Results showed that Mn increased interaction between LRRK2 and RAB10 assessed by proximity ligation assay (PLA) in the substantia nigra of WT mice, and this effect was exacerbated in G2019S (Fig. 4C), while Mn decreased interactions between 14-3-3ε and LRRK2 in the same region of WT, effects which were further decreased in LRRK2 G2019S mice (Fig. 4D). These results indicate that 14-3-3 proteins may play a role in regulating Mn-induced hyper LRRK2 kinase activity and cytotoxicity.

To gain more mechanistic insight, we also used an in vitro cell culture model of BV2 microglia. BV2 cells were transfected with human LRRK2 WT or G2019S expression vectors (Fig. 5A). Transfection with LRRK2 WT and G2019S vectors increased LRRK2 expression and phosphorylation at S1292 in WT-transfected BV2 cells, effects which were higher in G2019S cells (Fig. 5B). Mn exposure increased LRRK2 phosphorylation at S1292 in LRRK2 WT-BV2 microglia, which were further increased in G2019S-expressing BV2 cells (Fig. 5C). Mn also increased RAB10 phosphorylation in LRRK2 WT-expressing BV2 cells (Fig. 5C) and further increased in LRRK2 G2019S-expressing BV2 cells (Fig. 5D-E). Moreover, LRRK2 kinase inhibitors, MLi-2 and LRRK2-IN-1, which are well-established selective and highly specific in binding to LRRK2’s kinase domain and blocking its kinase activity (47,48), attenuated Mn-induced phosphorylation of LRRK2 and RAB10 in BV2 microglia expressing LRRK2 WT and G2019S, indicating that RAB10 is a downstream target substrate in Mn-LRRK2 activation (Fig. 5C-E).

**Figure 5.**
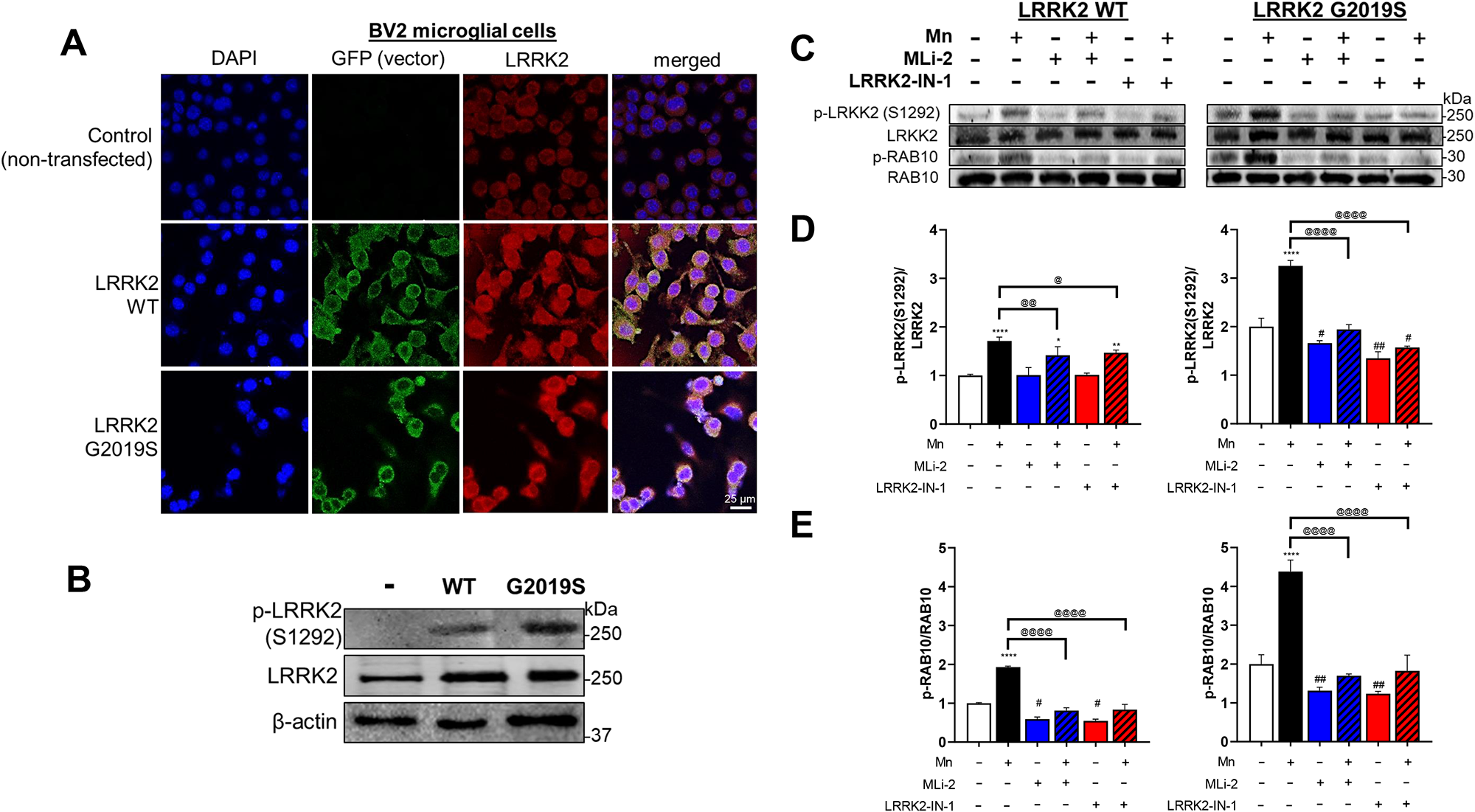
Mn further increases LRRK2 kinase activity in LRRK2 G2019S-expressing BV2 microglia cells. (A-B) Validation of transfection of vectors for LRRK2 WT and G2019S in BV2 microglia. (A) Fluorescence imaging of LRRK2 (red) in non-transfected control, LRRK2 WT-, and G2019S-expressing (green) BV2 cells. (B) Levels of LRRK2 phosphorylation (p-LRRK2, S1292) and protein in BV2 cells. Following LRRK2 WT and G2019S overexpression, LRRK2 inhibitors MLi-2 (50 nM, 0.5 h) and LRRK2-IN-1 (10 nM, 0.5 h) pre-treatment and Mn exposure, BV2 cells were analyzed for phosphorylation of LRRK2 at S1292 and RAB10 at T73 using western blotting. (C-E) Mn increased LRRK2 and RAB10 phosphorylation in LRRK2 WT BV2 cells, which were exacerbated in LRRK2 G2019S BV2 cells. Quantification of LRRK2 (D) and RAB10 (E) phosphorylation in BV2 cells. MLi-2 and LRRK2-IN-1 were used as LRRK2 inhibitors. β-actin was used as a loading control for protein. *p < 0.05, **p < 0.01, ****p < 0.0001, ^#^p < 0.05, ^##^p < 0.01, compared with the controls; ^@^p < 0.05, ^@@^p < 0.01, ^@@@@^p < 0.0001, compared with each other (two-way ANOVA followed by Tukey’s post hoc test; n = 3). Data are expressed as mean ± SD.

Next, given that microglia are the main neural cell types for regulating inflammatory reactions in the brain (16) and LRRK2 is implicated in Mn-induced microglial inflammatory responses (13), we tested if Mn-induced LRRK2 hyper kinase activity promotes inflammatory reactions in microglia. Studies have shown that Mn increased proinflammatory TNF-α in the mouse brain (43), and thus, we determined if Mn increases TNF-α in the striatum and midbrain of WT mice and if this effect is exacerbated in G2019S mice. Results showed that Mn increased TNF-α protein levels in the striatum and midbrain of WT mice, effects which were exacerbated in G2019S mice (Fig. 6A). These Mn’s in vivo effects were similarly exerted in BV2 cells expressing LRRK2 WT or G2019S (Fig. 6B-E). To determine if these Mn effects on TNF-α production are mediated by LRRK2 kinase activity, we tested these Mn effects in the presence of LRRK2 inhibitors, MLi-2 and LRRK2-IN-1. These inhibitors attenuated Mn-induced increase in TNF-α mRNA and protein levels in BV2 cells expressing LRRK2 WT or G2019S, indicating that TNF-α production is downstream effects of Mn-LRRK2 activation (Fig. 6B-E).

**Figure 6.**
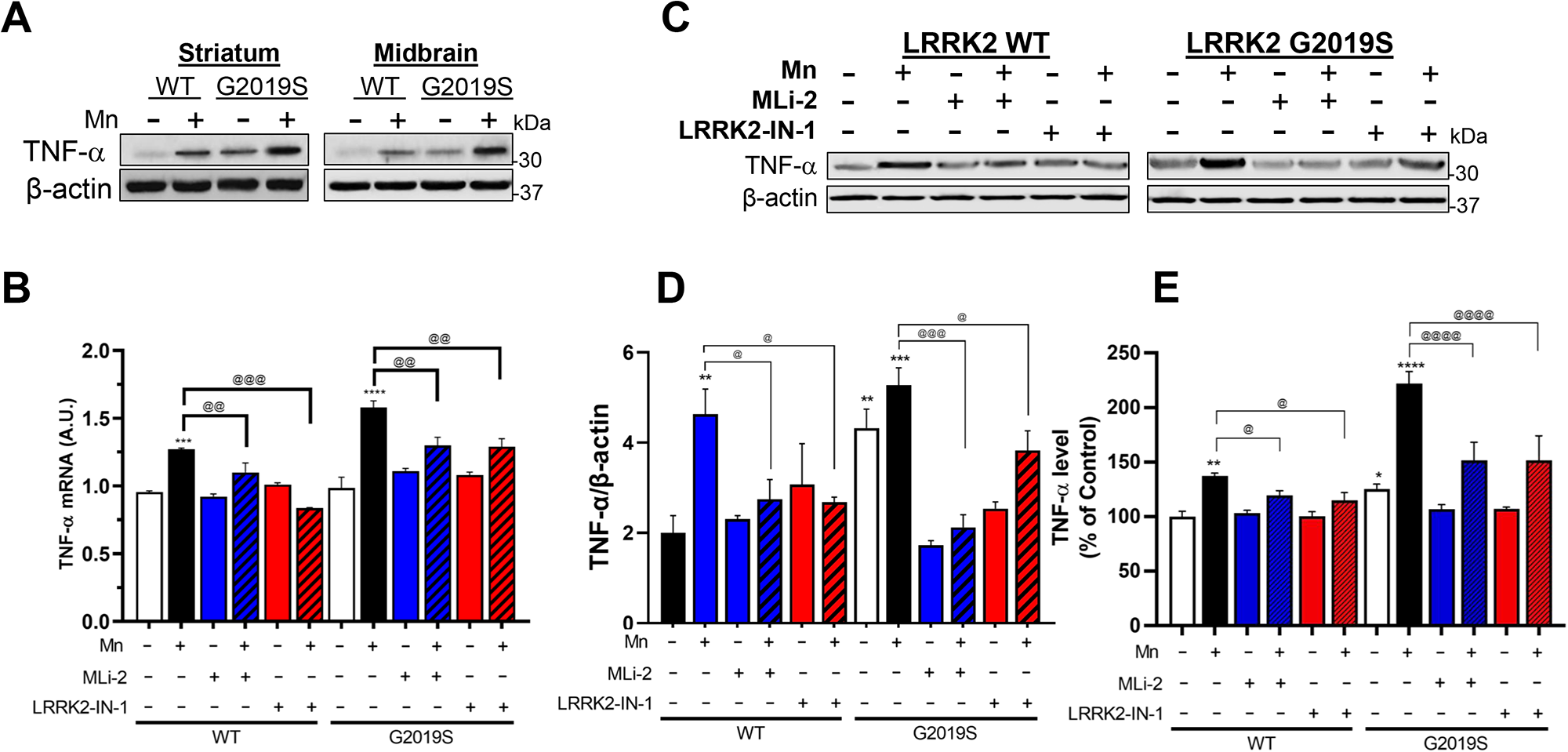
LRRK2 kinase activity plays a role in regulating Mn-induced proinflammatory TNF-α in mice and microglia. (A) After Mn exposure (MnCl_2_, 30 mg/kg, intranasal instillation, daily for 3 weeks), striatum and midbrain regions of mouse brains were analyzed for TNF-α protein levels by western blotting. (B-E) Following LRRK2 inhibitors MLi-2 (50 nM, 0.5 h) and LRRK2-IN-1 (10 nM, 0.5 h) pre-treatment and Mn exposure (250 µM, 12 h), LRRK2 WT- and G2019S-expressing BV2 cells were analyzed for TNF-α mRNA (B) and protein levels (C,D) by qPCR and western blotting, respectively. GAPDH and β-actin were used as normalization and loading control for RNA and protein, respectively. (E) Protein levels of secreted TNF-α were measured by ELISA. *p < 0.05, **p < 0.01, ***p < 0.001, ****p < 0.0001, compared with the controls; ^@^p < 0.05, ^@@^p < 0.01, ^@@@^p < 0.001, ^@@@@^p < 0.0001, compared with each other (two-way ANOVA followed by Tukey’s post hoc test; n = 3-5). Data are expressed as mean ± SD.

### LRRK2 mediates Mn-induced NLRP3 inflammasome activation and impairment of the autophagy-lysosomal pathway in mice and microglia

Mn activates the NLR family pyrin domain containing 3 (NLRP3) inflammasome in microglia and hippocampus of the mouse brain, leading to an inflammatory response (10). The NLRP3-IL-1β pathway is also implicated in PD pathogenesis (for review, see (49)). Thus, we tested if LRRK2 and G2019S modulate Mn-induced NLRP3 inflammasome activation. Results showed that Mn increased the primary protein components of NLRP3 inflammasome, such as NLRP3 and cleaved caspase-1 (CASP1), in the striatum and midbrain of WT mice, and this effect was exacerbated in G2019S mice (Fig. 7A). Lysosomal function was also assessed by determining the levels of lysosomal-associated membrane protein 1 (LAMP1) and active cathepsin B (CTSB) proteins as its dysfunction has been linked to NLRP3 inflammasome activation (50,51). Mn impaired lysosomal function in both the striatum and midbrain of the WT mice, as shown by decreasing LAMP1 and increasing CTSB protein levels, which were exacerbated in LRRK2 G2019S mice (Fig. 7A).

**Figure 7.**
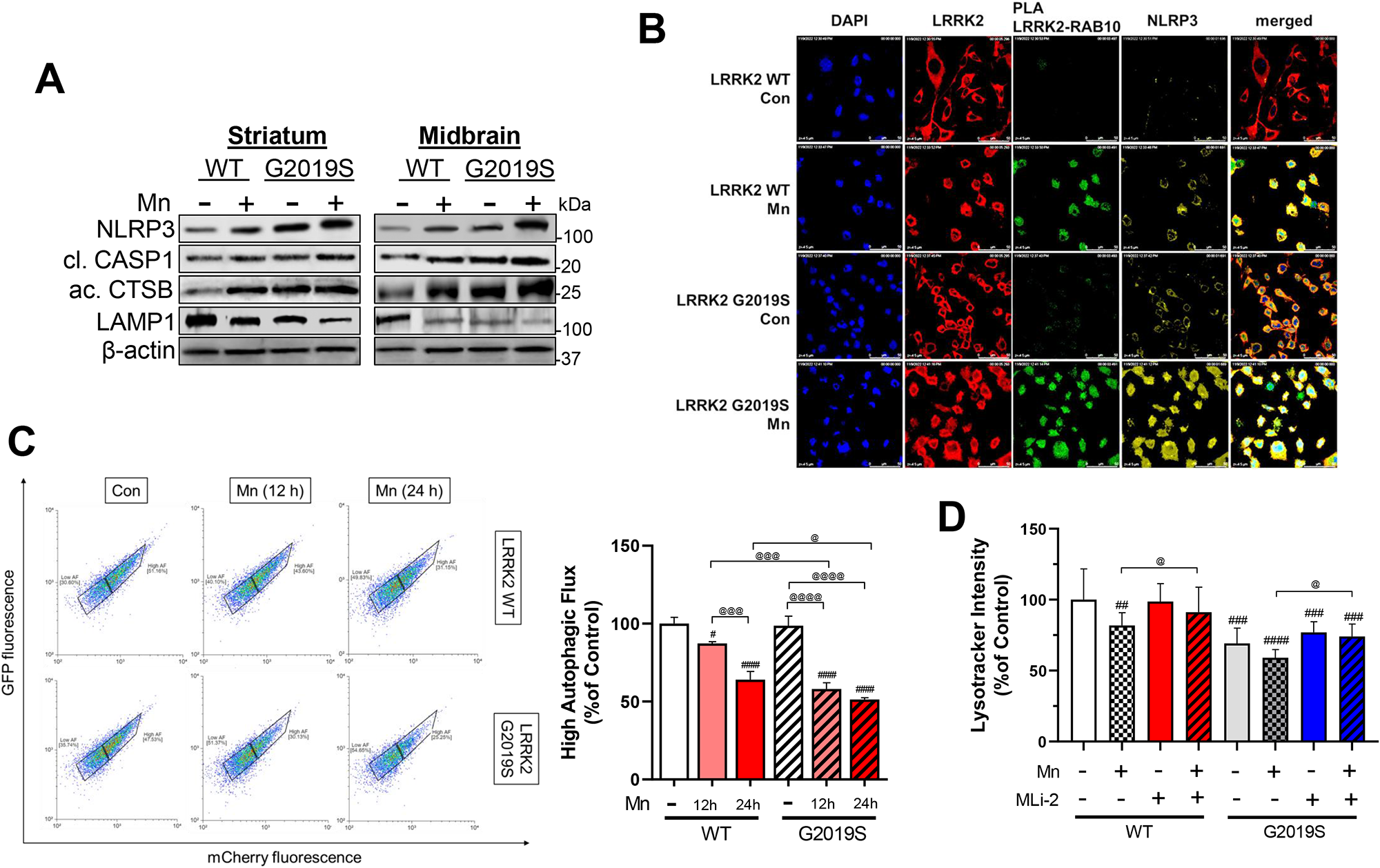
LRRK2 plays a role in Mn-induced NLRP3-IL-1β inflammasome pathway by modulating lysosomal function in mice and microglia. (A) After Mn exposure (MnCl2, 30 mg/kg, intranasal instillation, daily for 3 weeks), striatum and midbrain tissues were analyzed for the protein levels of NLRP3, cleaved CASP1, active CTSB, and LAMP1 by western blotting. (B) BV2 cells were examined for protein-protein interactions of LRRK2 with RAB10, fluorescence intensity, and colocalization of NLRP3 and LRRK2, RAB10 in the microglia. (C) Following Mn exposure for 12 and 24 h, LRRK2 WT and G2019S BV2 cells were assessed for autophagic flux with LC3-mcherry-GFP fluorescence assay. Cells with high autophagic flux were determined and quantified by red fluorescence using flow cytometry. (D) Following pre-treatment of LRRK2 inhibitor MLi-2 (50 nM, 0.5 h) and Mn exposure (250 µM, 12 h), LRRK2 WT and G2019S-expressing BV2 cells were analyzed for lysosomal activity by lysotracker assays. ^#^p < 0.05, ^##^p < 0.01, ^###^p < 0.001, ^####^p < 0.0001, compared with the controls; ^@^p < 0.05, ^@@^p < 0.01, ^@@@^p < 0.001, ^@@@@^p < 0.0001, compared with each other (two-way ANOVA followed by Tukey’s post hoc test; n = 3-5). Data are expressed as mean ± SD.

Next, we tested if LRRK2 is involved in Mn-induced increase of NLRP3 inflammasome activation as well as lysosomal impairment in BV2 cells expressing LRRK2 WT or G2019S. Results showed that Mn exposure and G2019S-expressing BV2 cells increased interaction between LRRK2 and RAB10, as reflected by increased PLA fluorescence. Mn also increased NLRP3 expression and its colocalization with LRRK2 in BV2 cells (Fig. 7B). G2019S-expressing BV2 cells increased NLRP3 expression compared to WT-expressing cells, which were further exacerbated by Mn (Fig. 7B). Similar to the in vivo data, Mn-induced lysosomal impairment, which likely contributed to NLRP3 inflammasome activation, is exacerbated in G2019S mice. In an in vitro model, BV2 cells were transfected with an autophagy reporter vector LC3-mCherry-GFP (52,53) in which red fluorescence indicates high autophagic flux activity. Results showed that Mn decreased number of cells exerting red fluorescence (Fig. 7C), indicating that Mn impaired autophagy activity in WT-expressing BV2 cells, which were exacerbated in G2019S-expressing BV2 cells (Fig. 7C). The lysotracker intensity analysis showed a parallel effect with Mn-induced autophagy impairment, while LRRK2 inhibitor MLi-2 abolished these Mn effects on autophagy, indicating that LRRK2 kinase activity is an upstream regulator of the Mn-induced autophagy impairment (Fig. 7D).

### Role of RAB10 in LRRK2-mediated Mn-induced NLRP3-IL1β pathway and lysosomal dysfunction in microglia

To determine the role of RAB10 as a LRRK2 target substrate in Mn-induced impairment of autophagy and inflammation, we assessed expression of p-RAB10 along with NLRP3 and CTSB in BV2 cells expressing WT and G2019S. The immunocytochemistry (ICC) imaging results showed that G2019S-expressing BV2 cells increased RAB10 phosphorylation, and G2019S-induced RAB10 activation was further enhanced by Mn (Fig. 8A). Mn increased fluorescence intensities of NLRP3 and CTSB along with higher colocalization with p-RAB10 (Fig. 8A). These Mn effects were further enhanced in G2019S-expressing BV2 cells. WB results showed that Mn increased protein levels of NLRP3, apoptosis-associated speck-like protein containing a CARD (ASC), cleaved CASP1, mature IL-1β, and active CTSB in WT-BV2 cells, which was exacerbated in G2019S cells (Fig. 8B, C). Furthermore, Mn-induced IL-1β release from WT-BV2 microglia was further exacerbated in G2019S-BV2 cells (Fig. 8D), while LRRK2 inhibitors MLi-2 and LRRK2-IN-1 abolished these effects in both WT- and G2019S-BV2 microglia (Fig. 8B-D), indicating that Mn-LRRK2 induces microglial inflammation, at least in part, by activating NLRP3 inflammasome.

**Figure 8.**
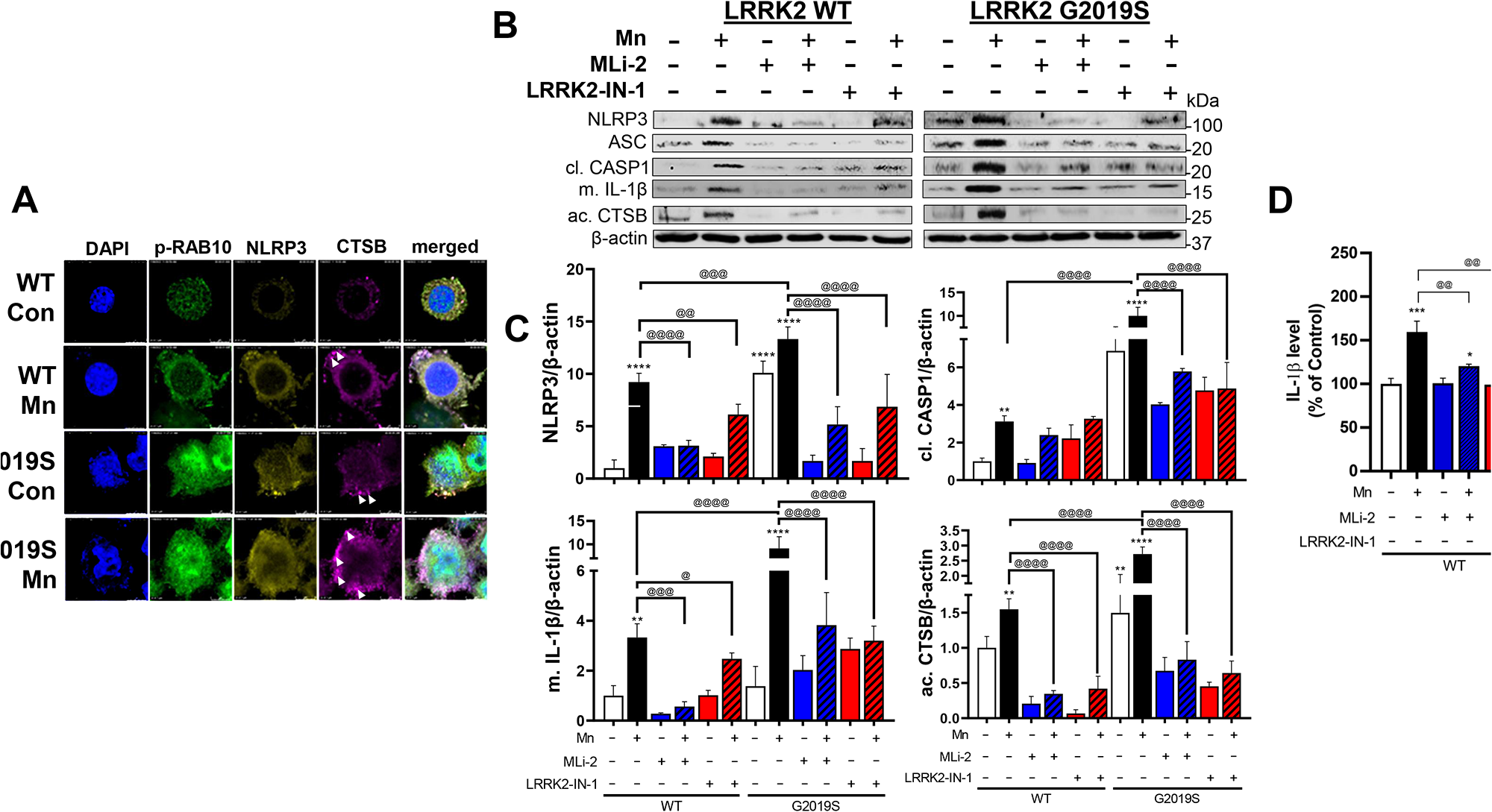
Mn increased NLRP3 inflammasome activation and proinflammatory IL-1β production via CTSB activity. (A-C) After transfection of LRRK2 WT and G2019S vectors following Mn exposure, BV2 cells were analyzed for the fluorescence intensity and colocalization of p-RAB10, NLRP3, and CTSB in microglia by immunofluorescence. (B) BV2 cells were analyzed for proteins by western blotting. (C) Protein levels of NLRP3, cleaved CASP1, mature IL-1β, and active CTSB were quantified in BV2 cells. β-actin was used as a loading control for protein. (D) Protein levels of secreted IL-1β in BV2 cell-free media were measured using ELISA. *p < 0.05, **p < 0.01, ***p < 0.001, ****p < 0.0001, compared with the controls; ^@^p < 0.05, ^@@^p < 0.01, ^@@@^p < 0.001, ^@@@@^p < 0.0001, compared with each other (two-way ANOVA followed by Tukey’s post hoc test; n = 3-5). Data are expressed as mean ± SD.

To further determine if RAB10 is involved in Mn-LRRK2-induced lysosomal dysfunction and NLRP3 inflammasome activation, we transfected BV2 cells with vectors of RAB10 expression and dominant negative-RAB10 (DN-RAB10) which represses RAB10. Transfection efficiencies of RAB10 and DN-RAB10 in BV2 cells were shown by GFP fluorescence (Fig. 9A). Results showed that Mn increased RAB10 phosphorylation in BV2 cells, which was decreased by DN-RAB10. Expression of RAB10 or DN-RAB10 did not affect Mn-induced LRRK2 phosphorylation in BV2 cells (Fig. 9A). In the absence of Mn, RAB10 expression showed higher p-RAB10 levels compared to DN-RAB10-expressing BV2 cells. Mn increased p-RAB10 levels in RAB10-expressing BV2 cells, whereas DN-RAB10 abolished this Mn effect (Fig. 9B). Mn also increased LRRK2 phosphorylation at S1292 to a similar extent in BV2 cells both with RAB10 and DN-RAB10 (Fig. 9B), indicating Mn-LRRK2 is the upstream regulator of RAB10. Given that RAB10 dysfunction has been shown to contribute to lysosomal defects and cytotoxicity (28), we determined if RAB10 function is involved in Mn-induced toxicity in BV2 cells. Results showed that Mn reduced cell viability in RAB10-expressing cells, which was blocked by DN-RAB10 and LRRK2 inhibitor MLi-2 in BV2 cells, which were parallel with alteration of lysosomal intensity (Fig. 9D). To determine the role of RAB10 in Mn-induced impairment of lysosomal function and NLRP3 inflammasome, we analyzed the protein levels of NLRP3, cleaved CASP1, LAMP1, active CTSB, and mature IL-1β. Mn increased NLRP3, cleaved CASP1, active CTSB, and mature IL-1β and reduced LAMP1 in RAB10-expressing cells, whereas these effects were attenuated by DN-RAB10 and MLi-2 (Fig. 9E). These data suggest that RAB10 is involved in Mn-induced NLRP3 inflammasomes via LRRK2 activity at least in part, by modulating lysosomal activity and CTSB in microglia.

**Figure 9.**
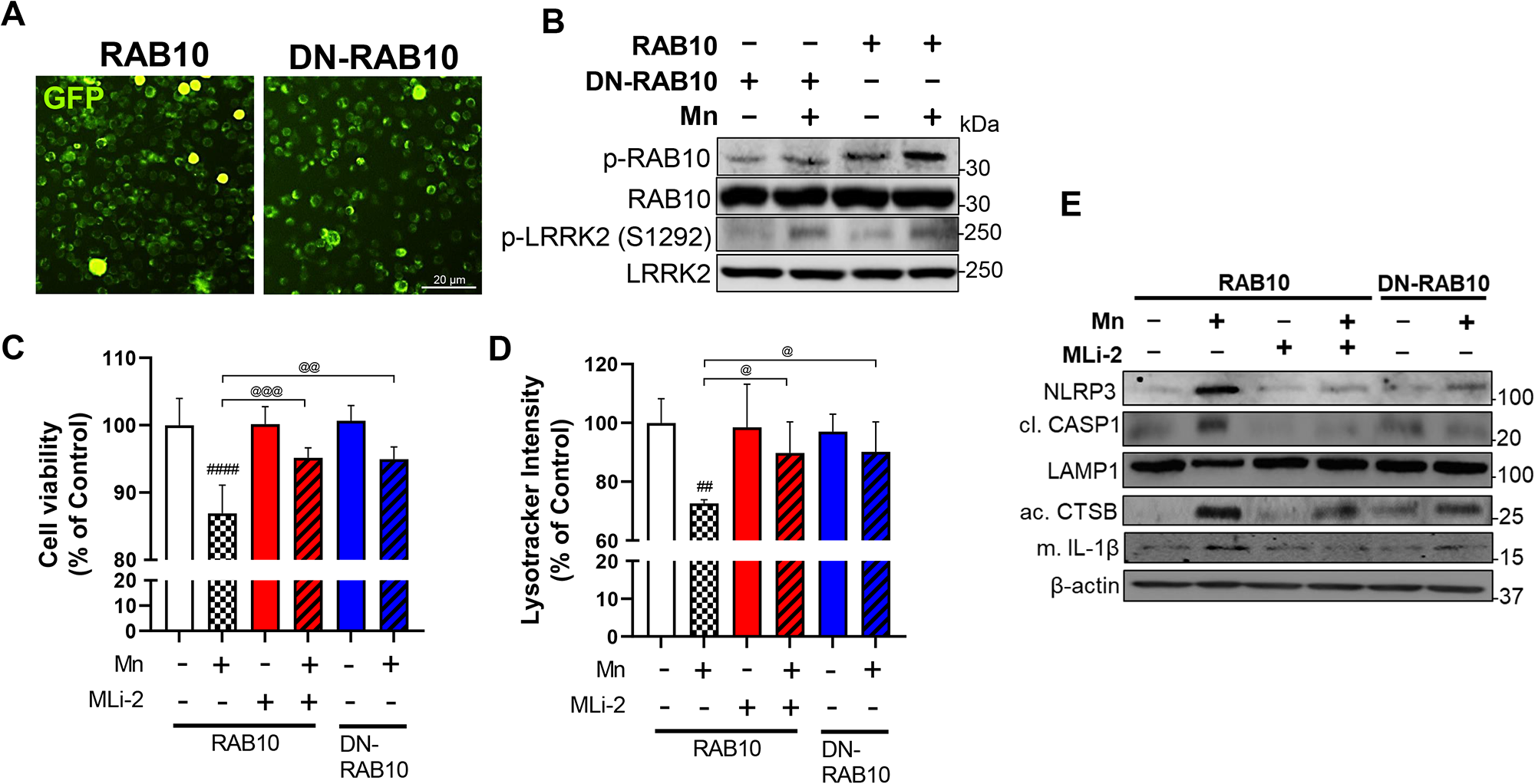
Mn alters lysosomal function in microglia via LRRK2-RAB10 activation. BV2 cells were transfected with RAB10 and DN-RAB10 vectors to modulate RAB10 function, followed by exposure to Mn (250 µM, 12h). (A) Validation of RAB10 and DN-RAB10 transfection in BV2 cells visualized with GFP fluorescence. (B-C) After Mn exposure, RAB10-and DN-RAB10-expressing BV2 cells were analyzed for proteins by western blotting. (B) Proteins for p-RAB10, RAB10, p- LRRK2, and LRRK2 were determined in RAB10- and DN-RAB10-expressing BV2 cells. (C-D) Following MLi-2 (50 nM, 0.5 h) pre-treatment and Mn exposure (250 µM, 12 h), RAB10- and DN-RAB10-expressing BV2 cells were assessed for cell viability (C) and lysosomal activity (D) and by resazurin and lysotracker assays, respectively. (E) Proteins for NLRP3, cleaved CASP1, LAMP1, active CTSB, and mature IL-1β were assessed in RAB10- and DN-RAB10-expressing BV2 cells after MLi-2 (50 nM, 0.5 h) pre-treatment and Mn exposure (250 µM, 12 h). ^##^p < 0.01, ^####^p < 0.0001, compared with the controls; ^@^p < 0.05, ^@@^p < 0.01, ^@@@^p < 0.001, compared with each other (two-way ANOVA followed by Tukey’s post hoc test; n = 5). Data are expressed as mean ± SD.

### Effect of Mn-induced microglial LRRK2 hyper kinase activity in mediating neuronal cytotoxicity

Next, we determined whether Mn-LRRK2-induced inflammation in microglia affects neighboring neuronal cells in an *in vitro* microglia-neuron co-culture system (Fig. 10A). Conditioned media (CM) from Mn-exposed BV2 cells expressing either LRRK2 WT or G2019S were applied to cath.a-differentiated (CAD) neuronal cells. CM was collected from BV2 cultures 6 h after Mn-containing media were replaced with fresh media to eliminate Mn’s direct effects on CAD cells. Results showed that treatment with CM from Mn-exposed BV2 cultures (LRRK2 WT+Mn-CM) increased neuronal cell toxicity as shown by decreased cell viability in CAD cells (Fig. 10B), which was further decreased by CM from Mn-treated G2019S-transfected BV2 cells (LRRK2 G2019S+Mn-CM). Moreover, CM from Mn-exposed BV2 microglia that were pre-treated with LRRK2 inhibitors (MLi-2 and LRRK2-IN-1) attenuated Mn-decreased cell viability in CAD cells (Fig. 10B). These results showed a similar pattern to reactive oxygen species (ROS) levels in BV2 cells (Fig. 10C, D).

**Figure 10.**
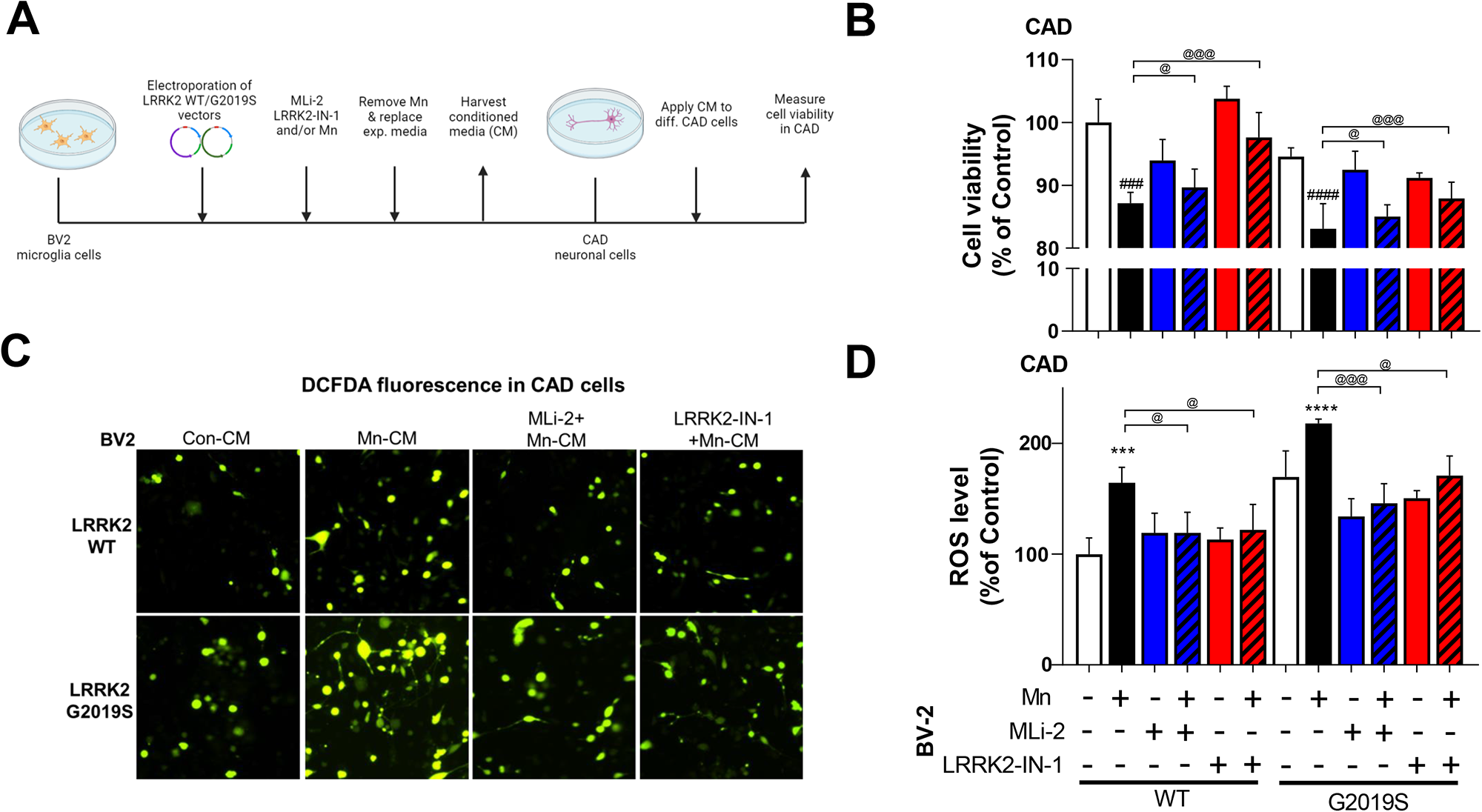
Microglial LRRK2 kinase activity mediates Mn-induced cytotoxicity in catecholaminergic neuron-like cells using a microglia-neuron co-culture. (A) BV2 cells were transfected with LRRK2 WT and G2019S vectors to modulate LRRK2 kinase activity, followed by LRRK2 inhibitors MLi-2 (50 nM, 0.5 h) and LRRK2-IN-1 (10 nM, 0.5 h) pre-treatment and Mn exposure (250 µM). After Mn exposure for 6h, BV2 experimental media were replaced with fresh media and incubated for additional 6 h prior to media collection as described in the Methods. These CM were applied to differentiated CAD cells. After CM exposure (12 h for cell viability; 3 h for ROS), CAD cells were analyzed for cell viability was determined by resazurin assays (B) and ROS levels by CM-H2DCFDA fluorescence (C, imaging; D, quantification). ***p < 0.001, ****p < 0.0001, ^###^p < 0.01, ^####^p < 0.001, compared with the controls; ^@^p < 0.05, ^@@@^p < 0.001, compared with each other (two-way ANOVA followed by Tukey’s post hoc test; n = 6). Data are expressed as mean ± SD.

### Mn-induced apoptosis was exacerbated in G2019S microglia

Given that LRRK2 is involved in Mn-induced cytotoxicity in microglia (13), we determined if hyper LRRK2 kinase activity plays a role in Mn-induced apoptosis in microglia by using LRRK2 inhibitors. Mn increased cytotoxicity in WT-BV2 cells, which was exacerbated in G2019S-BV2 cells, but inhibition of LRRK2 with MLi-2 or LRRK2-IN-1 attenuated Mn-induced reduction in cell viability (Fig. 11A). These toxic effects were parallel with Mn-induced modulation of apoptosis-related proteins, and were attenuated by LRRK2 inhibitors (Fig. 11B). It showed that Mn-induced modulation of Bcl-2, Bax, and active caspase-3, as well as the Bax/Bcl-2 ratio towards cell death in WT-BV2 cells were exacerbated in G2019S-BV2 microglia (Fig. 11C).

**Figure 11.**
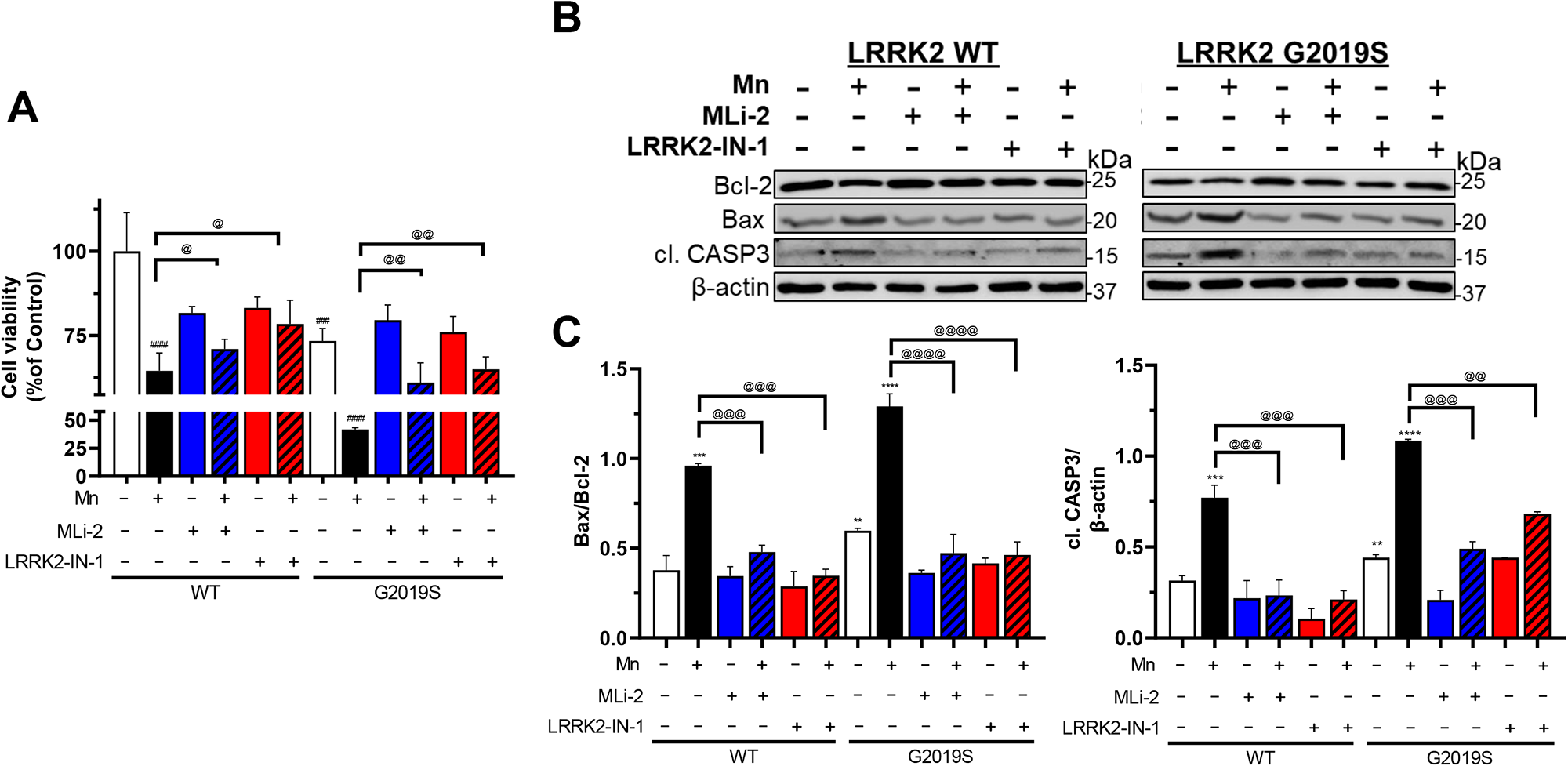
Mn-induced apoptosis is exacerbated by enhanced LRRK2 kinase activity in microglia. BV2 cells were transfected with LRRK2 WT and G2019S vectors to modulate LRRK2 kinase activity, followed by exposure to Mn (250 μM). (A) After treatment with LRRK2 inhibitors MLi-2 (50 nM, 0.5 h) and LRRK2-IN-1 (10 nM, 0.5 h) and Mn exposure (250 µM, 12 h) of BV2 cells for 24 h (for cell viability), cell viability was determined by resazurin assays . (B-C) After transfection with LRRK2 WT and G2019S, followed by LRRK2 inhibitors MLi-2 (50 nM, 0.5 h) and LRRK2-IN-1 (10 nM, 0.5 h) pre-treatment, and Mn exposure, protein levels for Bcl-2, Bax, active caspase-3 were analyzed by western blotting in BV2 cells. (C) Quantification of Bax/Bcl-2 ratio and active CASP3 in LRRK2 WT and G2019S were compared. β-actin was used as loading control for protein. **p < 0.01, ***p < 0.001, ****p < 0.0001, ^###^p < 0.001, ^####^p < 0.0001, compared with the controls; ^@^p < 0.05, ^@@^p < 0.01, ^@@@^p < 0.001, ^@@@@^p < 0.0001 compared with each other (two-way ANOVA followed by Tukey’s post hoc test; n = 3-6). Data are expressed as mean ± SD.

## Discussion

The present studies demonstrate that Mn exposure in WT mice induced motor deficits, cognitive impairment, and dopaminergic dysfunction in the nigrostriatal pathway with concomitant increase of LRRK2 and TNF-α protein levels in the basal ganglia, which were exacerbated in LRRK2 G2019S mice. In vitro studies with BV2 microglia transfected with LRRK2 WT and G2019S provided further insight to the role of LRRK2. Similar to the in vivo findings, Mn-induced proinflammatory TNF-α, IL-1β, and NLRP3 inflammasome activation in LRRK2 WT-expressing BV2 microglia were further exacerbated in G2019S-expressing BV2 cells. Moreover, Mn increased LRRK2 kinase activity as demonstrated by autophosphorylation of LRRK2 at S1292 and phosphorylation of RAB10, one of LRRK2’s substrates, in WT-microglia, effects which were more pronounced in G2019S-microglia. LRRK2 hyper kinase activity induced by Mn led to autophagy-lysosomal dysfunction, reflected by decreased LAMP1, and NLRP3 inflammasome activation and increased CTSB. Notably, LRRK2 inhibitors mitigated these Mn effects. These findings highlight the critical role of LRRK2 kinase in Mn toxicity in microglia and Mn’s contributing role in G2019S pathology via upregulation of LRRK2 kinase.

Given that Mn increased LRRK2 expression and its kinase activity in microglia (13), and LRRK2 G2019S mutation also increased LRRK2 kinase activity (34), LRRK2 kinase appears to be a common factor in Mn toxicity and G2019S pathogenesis. The Mn exposure treatment regimen used for the present study increased about two-fold of Mn levels in the striatum and midbrain (1), which is clinically relevant to human exposure levels (54). Since both Mn and G2019S are associated with dysfunction in the nigrostriatal pathway and an increase of LRRK2 kinase activity, our findings provide a link between Mn and LRRK2 G2019S pathology and critical insight into the potential role of LRRK2 kinase activity in Mn-induced neurotoxicity.

LRRK2 G2019S mutation in PD patients have been linked to both motor and cognitive impairments (40), and G2019S mutation impaired dopamine homeostasis in a mouse and C. elegans models (55,56). But G2019S mice did not show motor deficits and cognitive dysfunction in the present study likely because the mice were relatively young (8-weeks old) and LRRK2 mutation is associated with late-onset PD, presenting PD phenotype in later ages. The older G2019S mice (16 to 18 months old) exhibit PD pathology (55). Importantly, Mn exposure significantly reduced locomotor activity and memory function in young G2019S mice, suggesting that Mn is highly relevant environmental risk factor for G2019S-associated PD development since this mutation shows incomplete penetrance as gene-environment interaction. Although our studies showed that Mn increased LRRK2 expression in the striatum and midbrain regions, which contain the nigrostriatal dopaminergic neurons parallel with Mn-induced impairment of motor deficits and cognitive function, the role of LRRK2 in specific brain regions remains to be established since Mn levels are increased not only in the striatum and midbrain, but also other regions, such as hippocampus, cortex, and cerebellum (1). Moreover, G2019S mice were knocked in with LRRK2 G2019S in all tissues of the mouse, indicating that other cell types or tissues can indirectly modulate microglial G2019S-Mn toxicity. Cognitive impairment could be associated with multiple brain regions including cortex and hippocampus, requiring further research to determine regional-specific effects of Mn-LRRK2 pathology associated with behavioral deficits.

The Mn-induced reduction in interaction between LRRK2 and 14-3-3 protein were similar to studies showing that mutations in LRRK2 can disrupt its interaction with 14-3-3 proteins (57,58), leading to increased LRRK2 activity and toxicity, suggesting that 14-3-3 could be another molecular target in Mn-induced LRRK2 function. 14-3-3 proteins bind with LRRK2 in the cytoplasm, keeping LRRK2 inactive, which prevents LRRK2 hyperactivation and its toxic consequences (57,58). Multiple isoforms of 14-3-3 proteins, including γ, η, ζ, and ε, have been implicated in neurological disorders such as PD and schizophrenia (59). This study focused on the 14-3-3ε as it has shown protective effects against neurotoxins, rotenone, and MPP+ toxicity (60). Likewise, our finding of Mn-reduced interaction between 14-3-3ε and LRRK2 suggests a potential role of 14-3-3 in Mn-LRRK2 pathology.

Although the mechanism by which Mn increases LRRK2 expression has yet to be established, Mn increased LRRK2 proteins in the nigrostriatal pathway in both WT and G2019S mice, as well as colocalization of LRRK2 with Iba1, a marker of microglial activation in the substantia nigra, suggesting that microglial LRRK2 could play a critical role in Mn-induced dopaminergic neurotoxicity. As a result of Mn-increased LRRK2, proinflammatory cytokines and NLRP3 inflammasome-related proteins were produced in both *in vivo* and *in vitro* settings of WT and G2019S mice and microglia (Figs. 6, 8 and 10). Given the previous findings that Mn-induced these cytokines in microglia (14,15,43,61) and LRRK2’s inflammatory role in microglia (62,63), our current findings that LRRK2 inhibitors attenuated these Mn’s inflammatory effects shed further evidence on the critical role of LRRK2 in Mn’s inflammatory toxicity in microglia. Similar to previous reports that NLRP3 inflammasome increases mature IL-1β protein levels in microglia (10,64), Mn-increased IL-1β protein levels and release, at least in part by activation of NLRP3 inflammasome, which were exacerbated in G2019S cells (Fig. 7), possibly contributing to inflammation caused by Mn. Moreover, targeting LRRK2 activity may be a viable approach to mitigating Mn-induced NLRP3-mediated inflammatory toxicity.

Previous studies have reported that multiple RAB proteins such as RAB8a, RAB10, RAB, are LRRK2 substrates (65), and our findings that Mn activated RAB10 via LRRK2 as LRRK2 inhibitors blocked Mn-induced RAB10 phosphorylation, indicating that RAB10 plays a role in Mn-LRRK2 pathology. Interestingly, LRRK2 G2019S did not affect the total levels of RAB10 protein in microglia (Fig. 5C), suggesting that Mn and LRRK2 G2019S specifically affect RAB10 at the posttranslational level. Furthermore, since the LRRK2-RAB10 pathway is known to regulate functions of autophagy-lysosome and NLRP3 inflammasome (28), Mn-induced LRRK2 hyper kinase activity is likely to impair functions of lysosome and NLRP3 inflammasome in microglia. This Mn-induced activation of the LRRK2-RAB10 pathway may also impair autophagy function for removal of damaged organelles and proteins in Mn-induced toxicity. LRRK2 G2019S also showed elevated RAB10 activation compared to WT, underscoring the critical role of LRRK2-RAB10 in neurotoxicity and neurodegenerative disorders. LRRK2-induced optimal activation of RAB10 is essential for normal RAB10-mediated lysosomal function since LRRK2 hyper kinase activity could destabilize RAB10 in lysosomes, causing leakage and release of lysosomal enzymes such as CTSB (28,66). Studies have shown that CTSB interacted with the LRR domain of NLRP3 (51), and contributed to inflammasome activation (50). In addition, CTSB played a role in Mn-induced NLRP3 inflammasome in the hippocampus of mice (10). In line with these findings, our results showed that Mn increased colocalization of CTSB and NLRP3 in the cytoplasm of microglia, an effect which was more pronounced in G2019S. Although the involvement of LRRK2 in CTSB-induced NLRP3 activation cannot be determined from the present studies, RAB10 is involved in autophagy-lysosomal instability, thereby enhancing CTSB activity (Fig. 9). Moreover, both Mn and LRRK2 G2019S increased cytoplasmic CTSB, possibly by promoting lysosomal leakage since there were decreased LAMP1 protein levels and increased RAB10 phosphorylation in microglia.

Our findings also highlight the contribution of microglial LRRK2 in Mn-induced toxicity and subsequent neuronal damage. Elevated microglial LRRK2 kinase activity by Mn and G2019S mutation could exacerbate Mn-induced toxicity mechanisms, such as oxidative stress and apoptosis in adjacent neurons, corroborating previous reports that LRRK2 can mediate oxidative stress and apoptosis by altering proapoptotic and antiapoptotic protein levels (13). Further research is required to understand whether LRRK2 regulates Mn toxicity mechanisms via multiple signaling pathways and whether these pathways are inter-regulated or independent in microglia.

While our findings suggest Mn-induced toxicity in microglia via the LRRK2-RAB10-CTSB-NLRP3 pathway, a therapeutic strategy for mitigating Mn’s neurotoxicity may have some limitations as other dysregulated mechanisms, including mitochondrial dynamics, mitophagy, protein trafficking, and MAPK and NF-κB signaling may need to account for a full understanding on Mn-LRRK2 interactive mechanisms. Nonetheless, Mn-induced LRRK2 activation in microglia could be a potential target for treating Mn toxicity and other neurodegenerative diseases associated with elevated LRRK2 function. In addition, in the present study we tested only G2019S mutation; other LRRK2 mutations such as R1441G/C/H also showed elevated LRRK2 kinase activity may also be tested to determine the contributing role of Mn toxicity in other PD-related mutations. Moreover, exploring the in vivo effect of pharmacological inhibition of LRRK2 against Mn neurotoxicity and its potential side effects is crucial in determining the safety and efficacy of this potential therapeutic approach. To better understand Mn-LRRK2 pathology, further studies examining the molecular changes in neurons and astrocytes are necessary, as Mn appears to increase LRRK2 expression and kinase activity not only in microglia but also in other neural cell types (Fig. 4).

Taken together, our findings demonstrate that Mn-induced motor deficits and cognitive impairment were further exacerbated in G2019S mice, possibly via hyper kinase activity of LRRK2. Moreover, microglial LRRK2 appears to play a critical role in these Mn-LRRK2 toxicity, presenting a novel mechanism of Mn-induced neurotoxicity. Our findings underscore the role of LRRK2 in Mn’s neurotoxicity as gene-environment interaction. LRRK2-RAB10 played a critical role in Mn-induced autophagy-lysosomal dysregulation and NLRP3 inflammasome activation in microglia, leading to increase of CTSB levels, which is directly modulating NLRP3 inflammasome and adjacent neuronal health (Fig. 12). More research is needed to fully comprehend the molecular mechanism of LRRK2-RAB10 in Mn’s neurotoxicity. Nevertheless, the findings from the present study clearly indicate that LRRK2 hyper kinase is critically involved in Mn’s neurotoxicity and other neurodegenerative diseases associated with LRRK2 hyper kinase activity.

**Figure 12.**
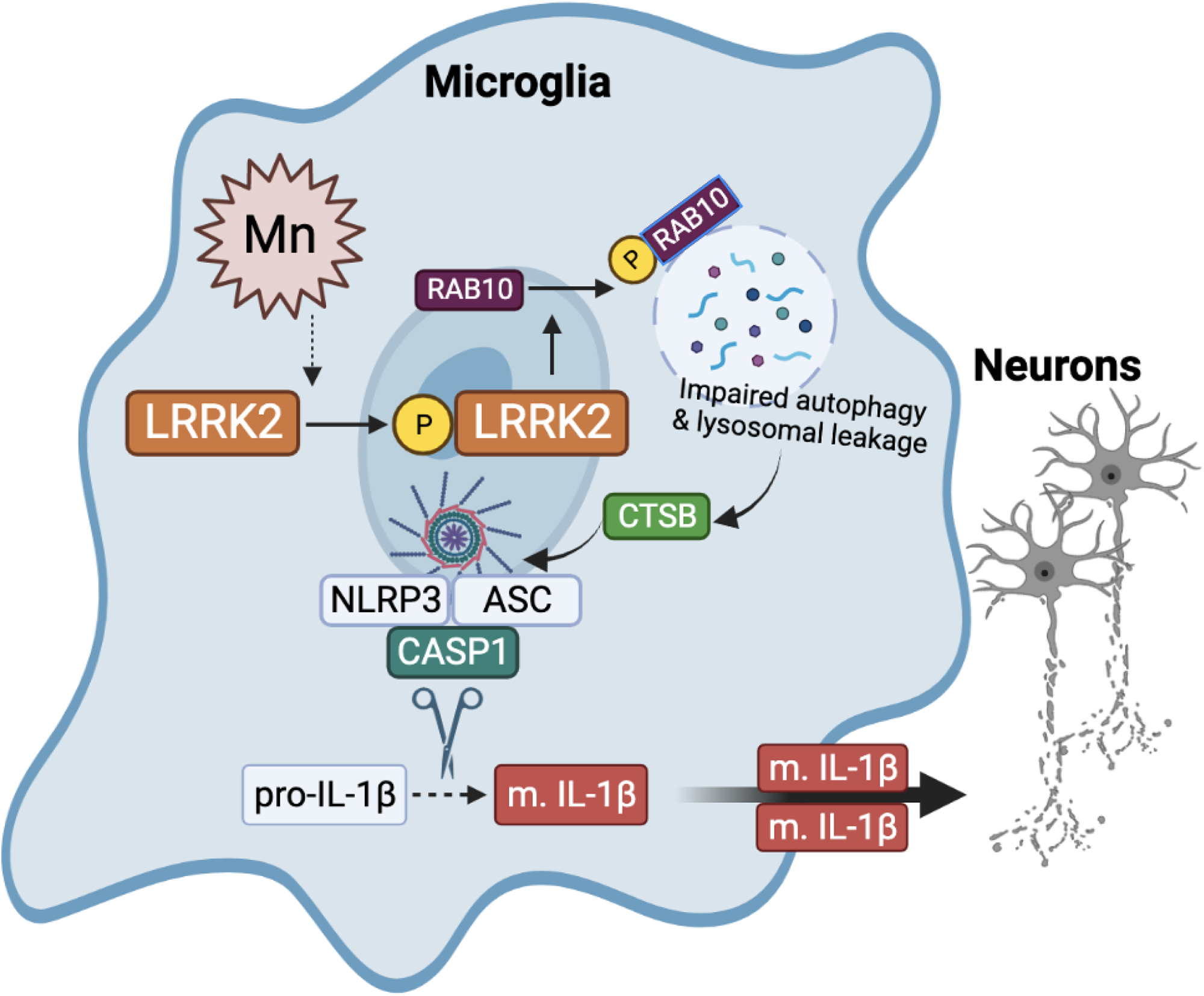
Proposed mechanism of LRRK2 kinase activity in Mn-induced NLRP3 inflammasome activation via RAB10 dysfunction in autophagy impairment and lysosomal enzyme leakage. Mn increases LRRK2 kinase activity, leading to autophosphorylation and phosphorylation of its target substrate, RAB10. RAB10 dysfunction contributes to Mn-induced autophagy impairment, reducing lysosomal membrane integrity and causing lysosomal cathepsin B (CTSB) leakage. The increased CTSB co-localizes and activates the NLRP3 inflammasome, resulting in the CASP1-mediated maturation of interleukin-1β (IL-1β) and subsequent neurotoxicity. This proposed mechanism provides insight into the potential role of LRRK2 kinase activity in Mn-induced neuroinflammation and highlights potential therapeutic targets for treating Mn-induced neurotoxicity.

### Experimental procedures

#### Chemicals, antibodies, and plasmid vectors

Manganese chloride (MnCl2), dimethyl sulfoxide (DMSO), LRRK2-IN-1 (438193), and resazurin sodium salt (R7017) were purchased from MilliporeSigma (St. Louis, MO). MLi-2 was purchased from R&D Systems (Minneapolis, MN). All cell culture media, including trypsin–EDTA and penicillin-streptomycin, Minimum Essential Media (MEM), Dulbecco’s Modified Eagle Medium (DMEM), and DMEM/F12 were obtained from Gibco (Carlsbad, CA). The chloromethyl derivative of 2’,’-dichlorodihydrofluorescein diacetate (CM-H2DCFDA, a ROS probe) and lysotracker deep red (L12492) were purchased from Invitrogen (Carlsbad, CA). Antibodies for TH (sc-25269), Iba1 (sc-28530), TNF-α (sc-52746), CTSB (sc-365558), ASC (sc-514414), IL-1β (sc-52012), CASP3,(sc-7272), Bcl-2 (sc-7382), Bax (sc-7480), LAMP1 (sc-20011), and β-actin (sc-47778) were obtained from Santa Cruz Biotechnology (Santa Cruz, CA). Antibodies for phosphorylated LRRK2 (S1292, ab203181), LRRK2 (ab133474) p-RAB10 (ab230261), RAB10 (ab104859), Iba1 (ab178846), NLRP3 (ab4207), rabbit anti-mouse (ab6728), donkey anti-goat (ab6885), goat anti-rabbit (ab97051) conjugated with horseradish peroxidase (HRP), and goat anti-rabbit, anti-mouse, anti-goat and anti-chicken antibodies conjugated with Alexa Fluor® 488, 568, or 647 were obtained from Abcam (Cambridge, MA). LRRK2 antibody (NB300-268) was from Novus Biologicals (Centennial, CO). Expression vectors for 2XMyc-LRRK2-WT (25361), 2XMyc-LRRK2-G2019S (25362), pDEST53-LRRK2-WT (25044), pDEST-LRRK2-G2019S (25045), EGFP-RAB10 (49472), EGFP-RAB10T23N (49545), and 2xmyc-RAB10 (164631) and reporter LC3-mCherry-GFP vector (110060) were from Addgene (Watertown, MA). The IL-1β and TNF-α standard tetramethylbenzidine (TMB) enzyme-linked immunosorbent assay (ELISA) development kit for mouse samples was acquired from Peprotech (Rocky Hill, NJ).

### Animals

All animal research protocols were approved by the Institutional Animal Care and Use Committee at Florida A&M University (FAMU) in Tallahassee, FL. Male C57BL/6 (WT, 8 weeks old) and LRRK2 G2019S knock-in (#13940-M, C57BL/6-*Lrrk2^tm4.1Arte^*, 8 weeks old) mice were purchased from Taconic Biosciences (Germantown, NY). Mice were housed in groups of five per cage and maintained on a 12-h light/dark cycle at a constant temperature of 22 ± 2 °C with ad libitum access to food, water, and enrichment. A total of forty male mice (20 LRRK2 WT and 20 LRRK2 G2019S) were used in this experiment. First, twenty mice were randomly assigned to four groups and were treated accordingly (n = 5): (1) WT plus vehicle, (2) WT plus Mn, (3) G2019S plus vehicle, and . (4) G2019S plus Mn. The mice were treated with Mn (MnCl_2_) at a dose of 30 mg/kg (330 µg of Mn, 1 µl per nostril in both nostrils) daily for 21 days, which is known to increase brain Mn levels twice to levels found in human brains with clinical Mn toxicity (54), as previously described in our studies (1). The remaining twenty mice were treated and grouped for acute Mn treatment (50 mg/kg, one time, i.p.), microdialysis, and dopamine measurement by high-performance liquid chromatography with an electrochemical detector (HPLC-ECD). Distilled water was used as a vehicle. Before and after Mn instillation, the mice were sedated with isoflurane for 3 min to prevent expulsion of the treatment from the nostrils.

### Open-field, rotarod, and NO recognition tests

Twenty-four hours after the last Mn treatment, an open-field test for locomotor activity and a rotarod test for motor coordination were conducted on the same day, as previously described (1). The open-field test was performed in a Plexiglas arena, and locomotor activity was measured using Fusion SuperFlex software v6.25 (Omnitech Electronics, Columbus, OH). Each mouse was acclimated to the arena for three consecutive days at the same time prior to the test day. Each mouse was placed in the center of the arena, and locomotor activity was measured for 30 min. Average walking distance, movement speed, and vertical activity were compared between groups.

The rotarod test was performed as described previously (1). Motor coordination was evaluated using the AccuRotor rotarod system (Omnitech Electronics). Mice were trained for three consecutive days, with each session consisting of three trials with 5 min breaks in between. During each trial, the mouse was placed on the rotating rod, and the speed was gradually increased up to 40 rpm by 0.1 revolution/sec over a 10 min period. The latency to fall was recorded using Fusion Software v6.3 for AccuRotor. The measurements of motor coordination were recorded for 650 sec in mice that persisted on the rod throughout each test. The average duration for each group was used for comparison.

For the NO recognition test, mice were first acclimated to the testing arena for three consecutive days. On the testing day, the mice underwent a familiarization phase, consisting of a 5 min break followed by a 10-min period in which they were exposed to two identical FO placed on the left back corner and right front corner of the open-field arena. During the NO recognition phase, one of the FOs was replaced with a NO that differed in texture, shape, and color, which was placed on the right front corner of the arena. The mice were then placed in the center of the arena and allowed to explore both FOs and the NO for 10 min. The time spent exploring the NO and FO was recorded using Fusion software. The time spent exploring the NO, and the discrimination index between the NO and FO were calculated and compared between groups (67). The average time spent exploring the NO and discrimination indices was used to compare groups.

### Microdialysis and measurement of striatal dopamine release

Mice were anesthetized with ketamine/xylazine (100 mg/kg) via intraperitoneal injection. Guide cannula (CXG-6, AMUZA, USA) and dummy cannula (CXD-6, AMUZA, USA) were implanted in the stereotaxic frame (68807, RWD, USA) according to the Paxinos mouse brain atlas, at a height, angle, and lateral position of mm: A +0.75, L −1.3, V −4.5 from Bregma (68). After one day of acclimatization, the dummy cannula was removed, and a microdialysis probe (CX-I-6-02, AMUZA, USA) was inserted into the guide cannula. Mice were acutely treated with Mn (single dose, 50 mg/kg, i.p.), followed by perfusion with artificial cerebral spinal fluid (aCSF) (147 mM NaCl, 2.8 mM KCl, 1.2 mM CaCl2, 1.2 mM MgCl2) at a flow rate of 1.5 µL/min (69) for 3 h. Fractions were collected every 20 min using a micro-fraction collector (FC-90, AMUZA, USA), with the first two fractions collected for baseline. All fractions were collected in tubes containing 5 µL of 0.2 M perchloric acid to prevent dopamine oxidation and stored at -80° C until analysis of the dialysate fraction. The flow rate was adjusted to 1.0 µL/min to stimulate dopamine release, and the dialysis fluid was switched to a solution containing 100 mM KCl 80 min after Mn treatment. After 20 min of K+ stimulation, the dialysis fluid was quickly switched back to standard aCSF (70). The collected fractions were analyzed using HPLC (HTEC-510, Eicom, USA) to measure dopamine levels. The fractions were then analyzed by the HPLC-ECD (HTEC-510, Eicom, USA) to measure extracellular dopamine levels in the striatum.

### Immunofluorescence and PLA

Immunohistochemistry (IHC) procedures were performed with minor modifications as previously described (1). Fixed and snap-frozen mouse brain tissue samples were processed to obtain coronal sections of substantia nigra (−2.70 to −3.40 mm) from the bregma, sliced into 30 µm thickness. IHC was conducted to assess the protein expressions of TH, LRRK2, and Iba1 in the substantia nigra, with tissue sections from three mice per group. Coronal sections on glass slides were washed twice with PBST (1X PBS, 0.3% Triton X-100) and incubated with blocking buffer (10% normal goat serum, 1% bovine serum albumin, and 0.3% Triton X-100 in 1X PBS) at room temperature for 1 h. The primary antibody solutions were incubated overnight at 4°C in a dark humidity chamber. Antibodies against LRRK2, TH, and Iba1 were diluted at 1:250 in the blocking buffer. The tissue sections were washed with a wash buffer, followed by a 2 h incubation at room temperature with secondary antibody solutions conjugated with Alexa Fluor 488, 568, and/or 647 fluorescent dyes diluted at room temperature 1:1,000. Slides were washed with wash buffer, dried, and mounted with a coverslip. The protein expressions from the same histological site of each sample were evaluated for fluorescence intensity using a Ts2R fluorescence microscope (Nikon Instruments, Melville, NY) and a Leica SPEII confocal microscope (Leica Microsystems Inc., Buffalo Grove, IL).

BV2 microglial cells were cultured on poly-L-lysine-coated circular coverslips for immunocytochemistry in 6-well plates. After appropriate treatment, cells were fixed with 4% paraformaldehyde in PBS pH 7.4 for 10 min at room temperature. The cells were then washed three times with ice-cold PBS and permeabilized with 0.1% Triton X-100 in PBS for 5 min. Subsequently, cells were incubated with a blocking buffer for 1 h at room temperature and then washed. The cells were then incubated overnight with primary antibodies for LRRK2, p-RAB10, RAB10, CTSB, or NLRP3 at 1:250 dilution at 4°C. After overnight incubation, cells were washed and incubated with fluorescent-conjugated secondary antibody Alexa Fluor® 488, 568, and/or 647 (1:1,000 dilution) for 1 h at room temperature. Finally, the cells were washed again and mounted on glass slides using DAPI-fluoromount solution for imaging analysis.

The PLA was performed according to the manufacturer’s instructions (MilliporeSigma). Briefly, coronal sections from tissue samples or cells fixed with 4% formaldehyde in PBS were prepared and blocked. Primary antibodies for LRRK2, RAB10, and/or 14-3-3ε (1:250 dilution) were incubated overnight at 4 °C. Oligonucleotide-conjugated secondary antibodies were then added and incubated at 37 °C for 1 h. The ligation solution containing two oligonucleotides and ligase was added and incubated at 37 °C for 30 min to hybridize two proximity oligonucleotides, which produces closed loops if the two interacting proteins are in proximity. An amplification solution was then added to enhance the fluorescent signals of hybridized oligonucleotides, and red fluorescent signals were visualized. For tissue sections, slides were washed and then mounted with a coverslip. For cells, relevant Alexa Fluor® conjugated secondary antibodies were added before final washing to determine the protein expression and cellular localization. Cellular localization and fluorescence intensity were assessed for each sample using a Ts2R fluorescence microscope (Nikon Instruments, Melville, NY) and a Leica SPEII confocal microscope (Leica Microsystems Inc., Buffalo Grove, IL).

### Cell culture and plasmid DNA transfection

Mouse BV2 microglia were grown in DMEM supplemented with 10% fetal bovine serum, 100 U/mL of penicillin, and 100 μg/mL of streptomycin. The cells were maintained at 37°C in a 95% air, 5% CO_2_ incubator. Mouse CAD catecholaminergic-like cells (MilliporeSigma) were grown in DMEM:F12 supplemented with 10% fetal bovine serum, 100 U/mL of penicillin, and 100 μg/mL of streptomycin, and 1% GlutaMAX. Growth media was replaced with a serum-free medium to differentiate CAD cells into morphologically and biochemically mature neurons.

For transfection of BV2 microglia, cells were transfected with the expression plasmids LRRK2 WT, LRRK2 G2019S, RAB10, and RAB10T23N by electroporation as previously described with slight modifications (71). Transfection involves introducing foreign genetic material (i.e., plasmid expression vector) into cells and is important in studying the function of specific genes. Cells were transfected with 10 µg of plasmid vector per 1.0 x 10^7^ cells. The electroporation parameters for exponential protocol were set at 200 V and 950-microfarad capacitance in a 4-mm electroporation cuvette. The transfected cells were then gently pipetted and incubated in a growth medium.

### Quantitative RT-PCR

After appropriate treatment, samples were prepared for quantitative PCR (qPCR). Total RNA was extracted from three samples per group using the RNeasy Mini Kit (Qiagen, Valencia, CA). Purified RNA (2 μg) was reverse-transcribed into cDNA using the high-capacity cDNA reverse transcription kit (Applied Biosystems, Foster City, CA). Real-time qPCR was performed on the CFX96 (Bio-Rad) using iQ SYBR Green Supermix (Bio-Rad, Hercules, CA) and 0.4 µM primers. The total reaction volume was 25 µL, with each cDNA template in 1 µL. The following primers were used for RNA extracted from mouse brain tissues and mouse BV2 microglia: mouse TNF-α, 5’-GAT GAG AAG TTC CCA AAT GGC-3’ (forward) and 5’-ACT TGG TGG TTT GCT ACG ACG-3’ (reverse); mouse GAPDH 5’-CTC ATG ACC ACA GTC CAT GC-3’ (forward) and 5’-CAC ATT GGG GGT AGG AAC AC-3’ (reverse). GAPDH was used as an internal control. The qPCR parameters were set for 1 cycle at 95°C for 10 min, 40 cycles at 95°C for 15 s, and 60-65°C for 1 min. Expression levels of each target gene were detected and quantified using the Bio-Rad CFX Manager version 3.1.

### Western blotting

Protein samples were collected from brain tissue and cell extracts for protein analysis by immunoblotting. After homogenization in a radioimmunoprecipitation assay buffer with protease inhibitors, the protein concentration was determined by a bicinchoninic acid assay. Equal amounts of protein were loaded onto 8 to 10% SDS-PAGE and analyzed by immunoblotting using antibodies at solutions ranging from 1:500 to 1:1000, followed by horseradish peroxidase-conjugated secondary antibody at a dilution of 1:5000. The blots were developed using the West Pico PLUS chemiluminescence substrate detection kit and imaged using the Bio-Rad ChemiDoc Imaging System. Protein levels were quantified using the Image Laboratory Software version 5.2.1 (Bio-Rad). Antibodies against p-LRRK2, LRRK2, p-RAB10, RAB10, Bcl-2, Bax, cleaved CASP3, TNF-α, NLRP3, ASC/PYCARD, cleaved CASP1, active CTSB, mature IL-1β, and LAMP1 were used to detect protein expression levels. β-actin was used as the loading control.

### Assays for cell viability, ROS, lysosomal activity, and autophagy flux

To investigate the effects of microglial LRRK2 on catecholaminergic/dopaminergic neuronal cells, differentiated CAD cells were treated with CM from Mn-exposed BV2 microglia expressing LRRK2 WT or G2019S, as described previously (71,72). BV2 microglia were transfected with LRRK2 WT and G2019S vectors and pre-treated with LRRK2 inhibitors MLi-2 and LRRK2-IN-1 before exposure to Mn. After Mn exposure for 6 h, the Mn-containing medium was replaced with a fresh experimental medium for an additional 6 h. The BV2-CM was then applied to CAD cells for ROS and cell viability assays. ROS was measured by incubating CAD cells with CM-H2DCFDA (ROS probe) for 30 min and measuring the green fluorochrome at 485/527 nm. Cell viability was measured by incubating cells with resazurin for 30 min at 37 °C and measuring the red fluorochrome from live cells at 530/590 nm. The endpoint intensities for green and red fluorochromes were detected using a Spectramax® i3x Multimode Microplate Reader (Molecular Devices, Sunnyvale, CA). The same cell viability assay was also used to determine Mn’s effects on the cell viability of BV2 microglia.

BV2 cells were transfected with LC3-mCherry-GFP, LRRK2 WT, and G2019S vectors to determine autophagy-lysosomal activity and incubated for at least 48 h before being treated with 250 µM of Mn for 12 or 24 h. Cells were dissociated with trypsin for 1 min, then deactivated with growth media, and harvested using centrifugation to obtain live cells. The cells were washed three times with DPBS at room temperature before undergoing flow cytometry analysis to quantify the fluorescence intensity of the LC3-mCherry-GFP reporter. Autolysosomes were quantified by counting mCherry-positive (red fluorescent) cells. Lysosomal activity was also measured by incubating BV2 cells with Lysotracker Deep Red for 30 min and measuring near infrared at 647/668 nm.

#### ELISA

To measure TNF-α and IL-1β release from BV2 cells, mouse ELISA kits (Peprotech) were used according to the manufacturer’s instructions. After exposure to Mn for the designated periods, a cell-free medium (1 mL/well) was collected for ELISA. The optical density of each well was measured using a Multimode Microplate Reader (Molecular Devices) set at 450 nm with wavelength correction set at 620 nm to determine the concentrations of secreted TNF-α and IL-1β.

### Statistical analysis

The results are expressed as the mean ± standard deviation (SD). Data analysis was conducted using GraphPad software version 9 (GraphPad, San Diego, CA). Bar graphs were used to visualize the differences in relative expression levels between groups. Statistical analysis was performed using two-way analysis of variance (ANOVA) due to two independent variables (genotypes and Mn treatment) for the open-field, rotarod, NO test, relative gene, and protein expression analyses, and cell toxicity assays. A p-value of less than 0.05 was considered statistically significant.

## Data Availability

All data are available within the article.

## Funding and additional information

This work was supported by the National Institute of Environmental Health Sciences, National Institutes of Health grants R01 ES024756 (to E. L.), R01 ES031282 (to E. L.), R01 ES10563 (to M. A.), R01 ES020852 (to M. A.), National Institute on Minority Health and Health Disparities, National Institutes of Health grant U54 MD007582 (to E.L.), and National Cancer Institute, National Institutes of Health grant SC1 CA200519 (to D.-S. S.). The content is solely the authors’ responsibility and does not necessarily represent the official views of the National Institutes of Health.

## Author Contributions

P. and E. L. conceptualization; E. P. and E. L. methodology; E. P. software; D.-S. S. and E. L. validation; E. P. formal analysis; E. P. investigation; E. L. resources; E. P., S.K., and A. D. data curation; E. P. and E. L. writing–original draft; E. P., S.K., A. D., M.D., D.-S. S., M. A., and E. L. writing–review and editing; E. P. visualization; E. L. supervision; E. L. project administration; D.-S. S., M. A., and E. L. funding acquisition.

## Conflict of Interest

The authors declare no conflicts of interest with the contents of this article.

## Abbreviations

The abbreviations used are:

aCSF: artificial cerebrospinal fluid
ATG5: autophagy-related 5
AD: Alzheimer’s disease; Bax
Bcl-2: associated X protein
Bcl-2: B-cell lymphoma 2
CASP1: caspase 1
CASP3: caspase 3
CM: conditioned media
CTSB: cathepsin B
cDNA: complementary DNA
DN-RAB10: dominant negative RAB10
ELISA: enzyme-linked immunosorbent assay
FO: familiar object
G2019S: glycine to serine substitution at amino acid 2019
GAPDH: glyceraldehyde 3-phosphate dehydrogenase
HMC3: human microglial cells 3
HPLC-ECD: high-performance liquid chromatography with electrochemical detector
HRP: horse-radish peroxidase
I2020T: isoleucine to threonine substitution at amino acid 2020
IHC: immunohistochemistry
IL-1β: interleukin 1 beta
KCl: potassium chloride
LAMP1: lysosomal- associated membrane protein 1
LRRK2: leucine-rich repeat kinase 2
MAPK: mitogen-activated protein kinase
NLRP3: nucleotide-binding domain, leucine-rich repeat-containing protein 3 or NOD-like receptor family, pyrin domain-containing 3
NO: novel object
PD: Parkinson’s disease
PBS: phosphate-buffered saline
PLA: proximity ligation assay
R1441C/G/H: arginine to cysteine/glycine/histidine substitution at amino acid 1441
RAB10: Ras-related protein Rab10; TNF-alpha, tumor necrosis factor-alpha
TH: tyrosine hydroxylase
WT: wild type

## Notes

### Competing Interest Statement

The authors have declared no competing interest.

### Summary of Updates

The abstract has been revised to clarify, and Figures 1-4 with relevant texts have been updated to clarify.

